# Cannabidiol promotes apoptosis and downregulation of oncogenic factors

**DOI:** 10.1101/2024.11.30.626177

**Authors:** Ashley Greenwood, Tomomi M. Yamamoto, Molishree Joshi, Kent Hutchison, Benjamin G. Bitler

## Abstract

Patients with high-grade serous carcinoma of tubo-ovarian origin (HGSC) often experience significant side effects related to their disease and treatments, such as pain, discomfort, nausea, and vomiting. Over the last two decades, the use of cannabinoids (CBD) to manage pain and anxiety has become more mainstream. However, there is limited data on how CBD interacts with HGSC tumor cells or whether CBD impacts the effect of chemotherapy.

Prior preclinical data has suggested the antitumor benefits of cannabinoids; however, the mechanism and data in ovarian cancer are limited. The objectives of this proposed research are to define the endocannabinoid system milieu in ovarian cancer, determine if CBD influences the growth of ovarian cancer cells, measure the cell viability when cannabinoids such as CBD are combined with standard-of-care therapies, and identify potential molecular pathways in which cannabinoids have a therapeutic effect. We conducted publicly available database searches, in vitro proliferation and apoptotic assays, functional protein signaling via reverse phase protein array analysis of CBD-treated cells using 2D cultured cells, and immunohistological analysis of ex vivo cultured patient-derived tumor slices treated with CBD.

Our data suggests that CBD is unlikely to affect the growth of cancer cells at physiologic doses but promotes apoptosis and can have growth inhibitory effects at higher concentrations. The inhibitory effects seen at high dose concentrations are likely from the upregulation of apoptotic pathways and inhibition of oncogenic pathways. Overall, physiologic CBD levels have minimal impact on cancer cell growth or chemotherapy efficacy.

## INTRODUCTION

Medicinal uses of cannabis sativa L. (marijuana) have origins dating back over 5000 years ago ^1^. Some historical medicinal properties documented include muscle spasms, vomiting, convulsions, rheumatism, tetanus, and rabies ^2^. However, in the modern era, the use of cannabinoids in medicine has been limited due to marijuana being categorized as a Schedule I controlled substance by the federal government ^3^. The Schedule I categorization has also led to a significant gap in cannabinoid biomedical research.

Over recent years, cannabis and its derivatives have been used for treating chemotherapy-induced nausea and vomiting, epilepsy, and multiple sclerosis, amongst other indications ^4^. Cannabinoids fall into three primary classifications that include 1) plant-based phytocannabinoids such as Δ^9^-tetrahydrocannabinol (Δ^9^-THC) and cannabidiol (CBD), 2) endocannabinoids that humans naturally produce, including anandamide (AEA) and 2- arachidonolyglycerol (2-AG),and 3) synthetic cannabinoids that have been manufactured to act similarly as the previously mentioned ^5^. CBD is a non-psychoactive phytocannabinoid that engages cannabinoid receptors such as the G Protein-Coupled Receptor cannabinoid receptor type-1(gene: *CNR1*; protein: CB1R) and cannabinoid receptor type-2 (gene: *CNR2*; protein: CB2R). Other CBD receptors have been identified such as transient receptor potential vanilloid type 1 (TRPV1), transient receptor potential vanilloid type 2 (TRPV2), transient receptor potential vanilloid type 3 (TRPV3), transient receptor potential vanilloid type 4 (TRPV4), serotonin receptor (HTR1A), Fatty acid amide hydrolase (FAAH), dopamine receptor (DRD2), G-protein-coupled receptor 55 (GPR55), transient receptor potential cation channel 8 (TRPM8), and peroxisome proliferator-activated receptor gamma (PPARy) ^6^. CBD is proposed to modulate these receptors, resulting in multiple therapeutic benefits like neuroprotection, antiepileptic, anxiolytic, antipsychotic, anti-inflammatory, analgesic, and anticancer properties ^7^.

There is increasing evidence that modulation of cannabinoid receptors can have tumor-suppressive properties in multiple tumor types. One of the earliest publications by Munson et al showed that orally administered Δ^9^-THC, Δ^8^-THC, and cannabinol inhibited tumor growth ^8^. THC and other cannabinoids have been shown to induce apoptotic death in glioma cells by CB1R/CB-dependent pathways by increasing the synthesis of ceramides. Ceramide activates the endoplasmic reticulum stress response, stimulating proapoptotic proteins, BAD, and BAX ^9^. While THC-induced apoptosis tends to be through a CB-dependent pathway, CBD exhibits activity through various non-GPRC mediated signaling as noted above. In a preclinical breast cancer model, CBD induced apoptosis through CB1R, CB2R, and vanilloid receptor-independent pathways ^10^. Greenhough et al. showed that THC induces apoptosis in colorectal cancers by CB1R-mediated inhibition of cancer survival pathways such as RAS/MAPK, ERK1, and PI3K/AKT^11^. Additionally, CBD, a partial agonist of CB1/CB2 receptors and antagonist of GPR55, may suppress mTOR/AKT signaling and activate proapoptotic proteins in colon cancer^12^.

There is a paucity of understanding on the impact of CBD on gynecologic malignancies. An article by Pineiro et al. discussed that the cannabinoid receptor, GPR55, modulates glysophospholipid lysophosphatidylinositol (LPI) in ovarian cancer cells. LPI induces calcium mobilization and activation of Akt and extracellular signal-regulated kinase (ERK)1/2 in these cells ^13^. In a study using immunocompromised mice, OVCAR5-derived established ectopic ovarian tumors treated with JWH-133, a CB2R agonist, showed an increase in endogenous cannabinoids AEA and 2-AG as well as a significant increase in tumor growth ^14^. In the clinical setting, there has been one case report of low-grade serous carcinoma having a complete resolution of disease following treatment with Laetrile in combination with CBD ^15^. There is a need to elucidate further the effect of CBD on ovarian cancer cells, especially in the more common and deadly high-grade serous carcinoma (HGSC) histotype.

In this study, we used publicly available datasets to understand the expression of patterns of CBD receptors. We investigated the effect of broad- and full-spectrum CBD on HGSC ovarian cancer cells. Full-spectrum CBD failed to inhibit the growth of HGSC cells. In contrast, broad-spectrum CBD effectively inhibited HGSC cell viability. Assessing the CBD-dependent signaling pathways, we found a significant enrichment in proteins associated with apoptosis pathways, including the PI3K/AKT/mTOR pathway. We confirmed that broad-spectrum CBD effectively induces apoptosis. Finally, in an *ex vivo* culture of a primary human HGSC tumor, there was a significant increase in apoptotic cells following CBD treatment.

## MATERIALS AND METHODS

### Cell Lines and Culture Conditions

Epithelial ovarian cancer cells OV7 [RRID: CVCL_2675] were cultured in Ham’s F12 media/ Dulbecco’ modified Eagle’s medium (DMEM) supplemented with 10% fetal bovine serum (FBS) 1% penicillin/streptomycin, 10ug/ml human insulin, 0.5ug/ml hydrocortisone. Epithelial ovarian cancer cells CAOV3 [RRID: CVCL_0201] were cultured in RPMI-1640 media supplemented with 10% FBS and 1% penicillin/streptomycin. Epithelial ovarian cancer cells TO14 [RRID:CVCL_2734] were cultured in RPMI-1640 media supplemented with 2mM Glutamine, 2mM Sodium Pyruvate, and 10% FBS. Cells were maintained at 37°C supplied with 5% CO2. Cell lines are used in culture for up to two months or 20 passages. Regular Mycoplasma testing was performed using LookOut Mycoplasma PCR detection (Sigma). Cells were last tested for Mycoplasma in April 2023. Cell lines were obtained from MilliporeSigma (10081203-1VL).

### Cannabinoid Products

Two cannabinoid products containing CBD were obtained from Ananada Professional. The broad-spectrum product contained 31.2mg/g cannabidiol with nondetectable levels of other cannabinoids. The hemp extract containing the CBD was suspended in hemp seed oil, glycerin, and placed in a gelatin-soft capsule. The oil was extracted from the gel capsule before use. The full spectrum CBD product also obtained from Ananada Professional contained a more diverse cannabinoid, including delta 9- tetrahydrocannabinol (0.63mg/g), delta 9 tetrahydrocannabinolic acid (0.50mg/g), CBD (11.86mg/g), cannabidiolic acid (20.39mg/g), cannabichoromene (0.62 mg/g), cannabichromenic acid (0.94mg/g), cannabigerol (0.35mg/g) and cannabigerolic acid (0.98mg/g). For in vitro assays, both the full and broad-sepctrum CBD were diluted and resuspended in a 1:1 Dimethylformamide (DMF) and medium-chain triglycerides (MCT) oil mixture. The diluted CBD products were vortexed on high for 30 seconds prior to use to create an emulsification.

### Colony formation assay

10,000 cells were seeded in 96 well plates and treated with increasing doses of broad and full spectrum cannabidiols up to 30μM (Ananada Professional). The cells were cultured for 72 hours. Colonies were fixed (10%methanol/10% acetic acid) and stained with 0.4% crystal violet. Crystal violet was dissolved in fixative and absorbance was measured at 570nm. Assays were performed in technical triplicate before reporting data.

### Incucyte Live Cell Imaging

OV7 and CAOV3 cell lines were seeded with 5000 cells in a 96-well plate. 24 hours later the cells were treated with increasing doses of cannabidiol doses and images with the Sartorius IncuCyte^®^ system for 72 hours to monitor cell proliferation percentage over time ^16^. Evaluation of apoptosis was evaluated in both cell lines using IncuCyte^®^ caspase-3/7 reagent (Sartorius, USA) that is non-fluorescent until activated by caspase-3/7 apoptosis activity. Once cleaved, the dye is capable of fluorescing green upon DNA binding, thus identifying the cells that underwent apoptosis. Incucyte imaging was supported by The University of Colorado Cell Technologies Shared Resource [RRID:SCR_023147)

### Reverse phase protein analysis

OV7 and CAOV3 cells were seeded in 10 cm culture dishes and treated with broad, full spectrum and combination cannabidiol. Cells were collected, snap frozen and sent to MD Anderson Functional Proteomics RPPA Core Facility (RRID:SCR_016649). Cell pellets were lysed and arrayed on nitrocellulose-coated slides. Sample spots were probed with 483 unique antibodies and visualized by DAB calorimetric reaction to produce stained slides. Relative protein levels were determined for each and designated as log2intensitiesand subsequently median-centered

### Gynecologic Tissue and Fluid Bank

The University of Colorado has an Institutional Review Board–approved protocol (COMIRB No. 07-935) to collect tissue from patients with both malignant and benign disease processes. All participants are counseled regarding the potential uses of their tissue. All patients have completed written informed consent per the Declaration of Helsinki and approved by the Colorado Multiple Institutional Review Board (COMIRB).

### Bioinformatic database analysis

The Cancer Genome Atlas (TCGA) Ovarian Serous PanCancer Atlas were accessed via the cBIOPortal (http://www.cbioportal.org) ^17 18^ HGSC tumor sc-RNA-seq data was analyzed via (https://lambrechtslab.sites.vib.be/^19^). Survival analyses (Kaplan–Meier curves), protein expression (*Z* score) analyses, and mRNA expression (transcripts per million) were examined.

### Statistical Consideration

Statistical analyses and P value calculations were performed using GraphPad Prism v10. Quantitative data are expressed as mean +/- standard error of the mean (SEM) unless otherwise noted. Analysis of variance (ANOVA) with Tukey multicomparison correction was used to identify significant differences in multiple comparisons. t-test was used for pairwise comparisons with False Discovery Rate method of Benjamini-Hochberg. Kaplan-Meier and Logrank was used for survival analysis. Dose response curves were analyzed via non-linear regression followed by a comparison of IC50 using the Extra Sum of Squares F Test. All experiments were completed in three independent experiments and at minimum of triplicate. For all statistical analyses, the level of significance was set at 0.05.

### Data availability statement

Data are available upon request from the corresponding author.

## RESULTS

### Profiling the CBD system in ovarian cancer

There is a plethora of receptors that are known to interact with CBD, and little is known about their expression in HGSC tumors. Using the TCGA dataset for HGSC, we investigated the mRNA expression of 12 predicted CBD receptors. In 377 HGSC tumors, TRPV2 (TPM+1=6.311, SD=4.246), TRPV4 (TPM+1=6.518, SD=7.729), PPARG (TPM+1=1.998, SD=3.238), and FAAH (TPM+1=17.63, SD=9.553) had the highest expression across the 12 receptors (**Figure 1A**). Assessing the same 12 receptors across 65 HGSC cell lines, a similar pattern was observed, with TRPV2 (TPM+1=1.069, SD=1.366), PPARG (TPM+1=2.506, SD=2.074), and FAAH (TPM+1=2.387, SD=1.329) having the highest expression (**Figure 1B**).

**Figure 1.**
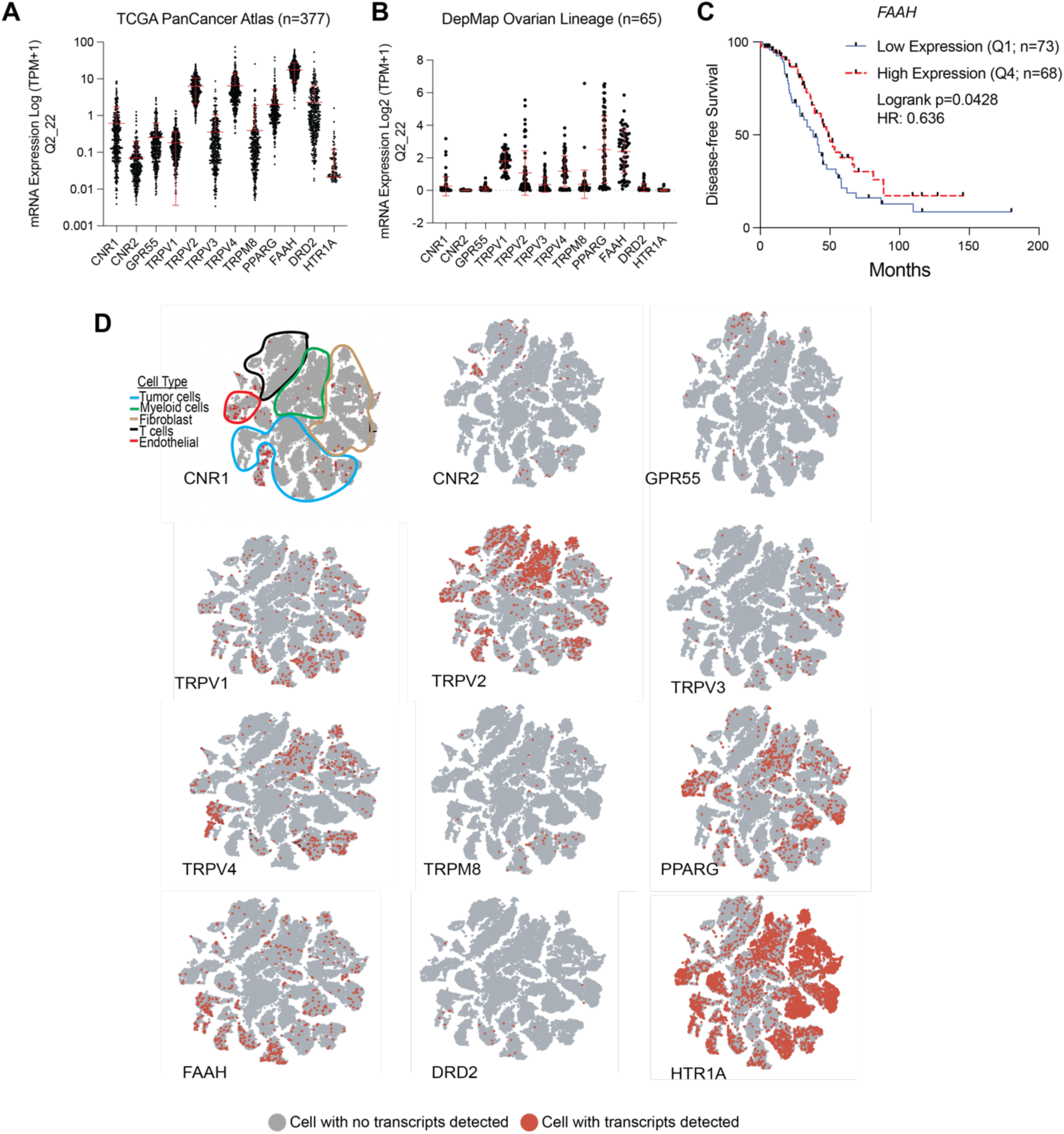
Evaluation of the endocannabinoid system in HGSC tumors and ovarian cancer cells. **A)** mRNA expression of predicted receptors/effectors of the endocannabinoid receptors in the HGSC tumors (PanCancer Atlas, TCGA; n=377). **B.** mRNA expression of predicted receptors/effectors of the endocannabinoid receptors in the HGSC cell lines (DepMap; n=65). C) Disease-free survival of patients with HGSC tumors with low (1^st^ Quartile) or high (4^th^ Quartile) *FAAH* expression. **D)** mRNA expression of predicted receptors/effectors of the endocannabinoid receptors in five HGSC tumors used for single-cell RNA-sequencing ^19^. Red dots indicate the detection of expression. Error bars, SD.

We next assessed the prognostic value and dependency of the expression of each of the receptors; using the TCGA HGSC tumor and DepMap expression data, respectively. We determined the groups based on LOW (Quartile 1) and HIGH (Quartile 4) mRNA expression. Across all 12 receptors, only elevated expression of FAAH correlated to a significant disease-specific survival benefit (p=0.0428, q=0.171, Hazard Ratio: 0.636, 95% CI 0.410-0.987) (**Table 1** and **Figure 1C**). In 60 HGSC cell lines, the CRISPR knockout of each individual receptor did not significantly impact cell viability based on the Chronos Dependency Score (**Table 1**). However, FAAH knockout demonstrated the lowest Chronos Dependency Score (Avg: −0.204, SD: 0.127), suggesting that loss of FAAH reduced cell viability in HGSC cell lines.

**Table 1.** Survival Analysis of predicted receptors/effectors. PanCancer Atlas. Analysis was conducted using low (Quartile 1) and high (Quartile 4) mRNA expression for each indicated gene. Logrank p-values are indicated.

| GENE | Disease Specific Survival (p-value) | Progression Free Survival (p-value) | Overall Survival (p-value) | Disease Free Survival (p-value) | CRISPR Dependency Score (CHRONOS) | Chronos Std. Dev. |
| --- | --- | --- | --- | --- | --- | --- |
| CNR1 | 0.992 | 0.332 | 0.627 | 0.246 | 0.035 | 0.093 |
| CNR2 | 0.436 | 0.73 | 0.524 | 0.573 | 0.021 | 0.101 |
| GPR55 | 0.763 | 0.402 | 0.808 | 0.382 | 0.174 | 0.110 |
| TRPV1 | 0.875 | 0.823 | 0.962 | 0.962 | 0.011 | 0.119 |
| TRPV2 | 0.48 | 0.214 | 0.569 | 0.448 | -0.072 | 0.091 |
| TRPV3 | 0.833 | 0.593 | 0.484 | 0.853 | 0.130 | 0.098 |
| TRPV4 | 0.069 | 0.144 | 0.071 | 0.17 | 0.040 | 0.100 |
| TRPM8 | 0.133 | 0.095 | 0.285 | 0.686 | -0.101 | 0.097 |
| PPARG | 0.992 | 0.862 | 0.896 | 0.171 | 0.000 | 0.121 |
| FAAH | **0.043** | 0.39 | 0.088 | 0.546 | -0.204 | 0.127 |
| DRD2 | 0.359 | 0.39 | 0.682 | 0.774 | 0.128 | 0.102 |
| HTR1A | N/A | N/A | N/A | N/A | -0.056 | 0.107 |
| CNR1+CNR2 | 0.431 | 0.958 | 0.262 | 0.259 |  |  |
| FAAH+CNR1 | 0.15 | 0.126 | 0.556 | 0.123 |  |  |
| FAAH+CNR2 | 0.375 | 0.553 | 0.422 | 0.485 |  |  |
| TRPV3/TRPV4 | 0.193 | 0.202 | 0.142 | 0.612 |  |  |

A limitation of the TCGA expression dataset is that it is challenging to define gene expression patterns within specific cell types in the tumor microenvironment. To overcome this limitation, we extended the profiling of the 12 CBD receptors into a single-cell RNA-sequencing (scRNA-seq) dataset from five primary HGSC tumors^19^. In this analysis, we were only interested in whether specific cell types expressed the designated receptor. Prior research has primarily focused on CBD receptors 1 and 2 (CNR1/2), and we observed that most cell types actually had few cells that had detectable transcripts of CNR1 and CNR2, while HTR1A was expressed broadly throughout most of the cell types (**Figure 1D**). Regarding cell type-specific expression, FAAH expression was mainly restricted to tumor cells, and TRPV2 expression was predominantly detected in myeloid cells and T cells. Taken together, these data highlight that the components of the CBD system are intact within the HGSC tumor compartment, have limited prognostic value, and are expressed in tumor cells.

### Establish the CBD-dependent effects on cell viability alone and with therapeutic agents

Next, we wanted to determine the growth effect of CBD on ovarian cancer cell lines. We selected the OV7 and CaOV3 cell lines based on their determination to be serous carcinoma histotypes and expression of multiple CBD receptor transcripts^20^. After being treated with a dose escalation of broad- and full-spectrum CBD, we performed colony formation assay as a measure of cell viability in OV7 and CAOV3 cells. Compared to vehicle control (DMF/MCT), we observed a dose-dependent reduction in crystal violet absorbance in both cell lines. In both cell lines, the full-spectrum CBD had minimal impact on colony formation. In the OV7 cell line, the 6 μM broad-spectrum reduced cell viability, and the 30 μM of broad-spectrum CBD significantly reduced cell viability (**Figure 2A**). In the CaOV3 cell line, both 6 and 30 μM of broad-spectrum CBD significantly reduced colony formation (**Figure 2B**). These data suggest mechanistic differences in the activity of full and broad-spectrum CBD.

**Figure 2.**
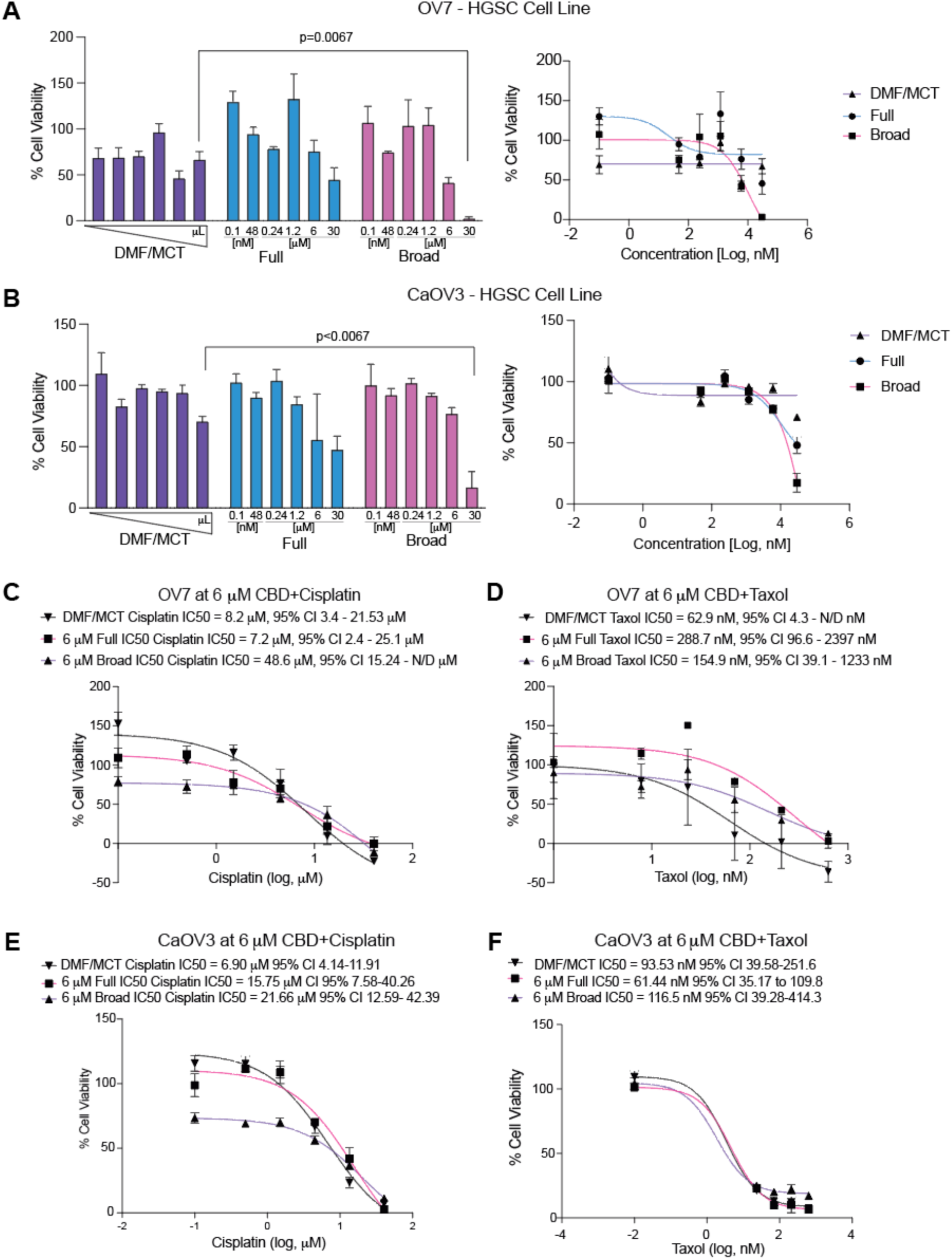
Broad-spectrum CBD inhibits cell proliferation and does not antagonize the response to standard-of-care chemotherapy. **A)** OV7 and **B)** CaOV3 HGSC cell lines were treated with increasing doses of full-spectrum and broad-spectrum CBD. After 72 hours, colony formation followed by crystal violet staining was used to assess a dose-response to CBD and cell viability. **C)** OV7 cells were treated with a combination of 6 mM Full or Broad-spectrum CBD with increasing doses of cisplatin or **D)** taxol. Dose-response curves and 50% maximal inhibitory concentration (IC50) were calculated. **E)** CaOV3 cells were treated with the combination of 6 mM Full or Broad-spectrum CBD with increasing doses of cisplatin or **F)** taxol. Dose-response curves and 50% maximal inhibitory concentration (IC50) were calculated. Error bars, SD. Statistical test, multicomparison ANOVA with Tukey correction.

We next wanted to assess the effect of broad- and full-spectrum CBD on the response to standard-of-care chemotherapy agents, cisplatin and taxol. Using a similar colony formation assay, we treated cells with a standard 6 μM CBD dose and then with increasing doses of chemotherapy (Cisplatin, 0.5 to 40 μM or Taxol, 23.3 to 630 nM). Following 72 hrs, we calculated the 50% maximal inhibitory concentration (IC50) to determine the effect of CBD.

In the OV7 cells treated with 6 μM CBD and cisplatin, compared to DMF/MCT (IC50 = 8.2 μM, 95% CI 3.4-21.53 μM), we observed that full spectrum did not shift the IC50 (IC50 = 7.2 μM, 95% CI 2.4 - 25.1 μM). In contrast, the broad Spectrum CBD did increase the cisplatin IC50 (IC50 = 48.6 μM, 95% CI 15.24 μM - N/D) but was not statistically different than DMF/MCT and had overlapping confidence intervals (**Figure 2C**). In the OV7 cells when treated with 6 μM CBD and taxol both Full (288.7 nM 95% CI 96.6 nM-2397 nM) and Broad Spectrum (154.9 nM 95% CI 39.1-1233 nM) CBD led to an increase IC50, compared to DMF/MCT (62.9 nM 95% CI 4.3 nM-N/D), however, the shifts were not significant based on overlapping confidence intervals (**Figure 2D**).

In the CaOV3 cells treated with 6 μM CBD and cisplatin, compared to DMF/MCT (6.9 μM, 95% CI 4.1 μM - 11.91 μM), both Full (15.75 μM, 95% CI 7.58 μM - 40.26 μM) and Broad Spectrum CBD (21.66 μM, 95% CI 12.59 μM - 42.39 μM) led to a shift in the IC50, suggesting that CBD and cisplatin may be antagonistic (**Figure 2E**). In the CaOV3 cells when treated with 6 μM CBD and taxol, when compared to DMF/MCT (93.53 nM, 95% CI 39.58 - 251.6 nM), neither Full (61.44 nM, 95% CI 35.17 nM - 109.8 nM) or Broad Spectrum (116.5 nM, 95% CI 39.28-414.3 nM) CBD significantly shifted the IC50 of taxol (**Figure 2F**). This data suggests CBD has minimal interaction with chemotherapies and the possible limited cisplatin:CBD interaction is cell-line dependent.

### Functional protein signaling assays to assess CBD-mediated signaling

To identify changes in signaling and protein expression, OV7 and CAOV3 cells were treated with 6 μM of broad-spectrum CBD for 72 hours and analyzed via reverse phase protein array (RPPA). Based on the prior studies, it was anticipated that the decrease in cell viability was identified with the colony formation analysis due to apoptosis and PI3K/AKT/MTOR signaling ^21,22^. In the OV7 cells, 54 proteins were significantly differentially regulated (**Figure 3A and Table S1**). In the CAOV3 cells, 27 proteins were significantly differentially regulated (**Figure 3B and Table S1**). Using immunoblotting, we confirmed that 6 μM Broad spectrum CBD reduced the expression of phosphorylated CDK1 (T14) and ZEB1 (**Figure 3C-D**) in the OV7 cell line. The proteins were then analyzed via Gene Set Enrichment to identify overlaps with Hallmark Pathways. While there was no overlap in differentially regulated proteins between the CAOV3 and OV7 cells, the affected pathways identified between the two cell lines following CBD treatment significantly overlapped (**Figure 3E**). The CAOV3 identified 18 significant (adj. p<0.05) pathways, while OV7 had 23, and there was an overlap in 13 pathways. The overlap represents a 1.57-fold enrichment (p=0.006). Assessing the Hallmark Pathways identified in the overlap between the two cell types, the most significant changes were seen with apoptosis and PI3K/AKT/MTOR signaling (**Figure 3F**). Other notable pathways included the Reactive Oxygen Species pathway, which is an important aspect of chemotherapy response ^23^. Taken together, broad-spectrum CBD modulates protein signaling. However, given the lack of specific overlap in differentially regulated proteins, the mechanism of CBD-dependent signaling is potentially due to more broad effects, such as the induction of reactive oxygen species.

**Figure 3.**
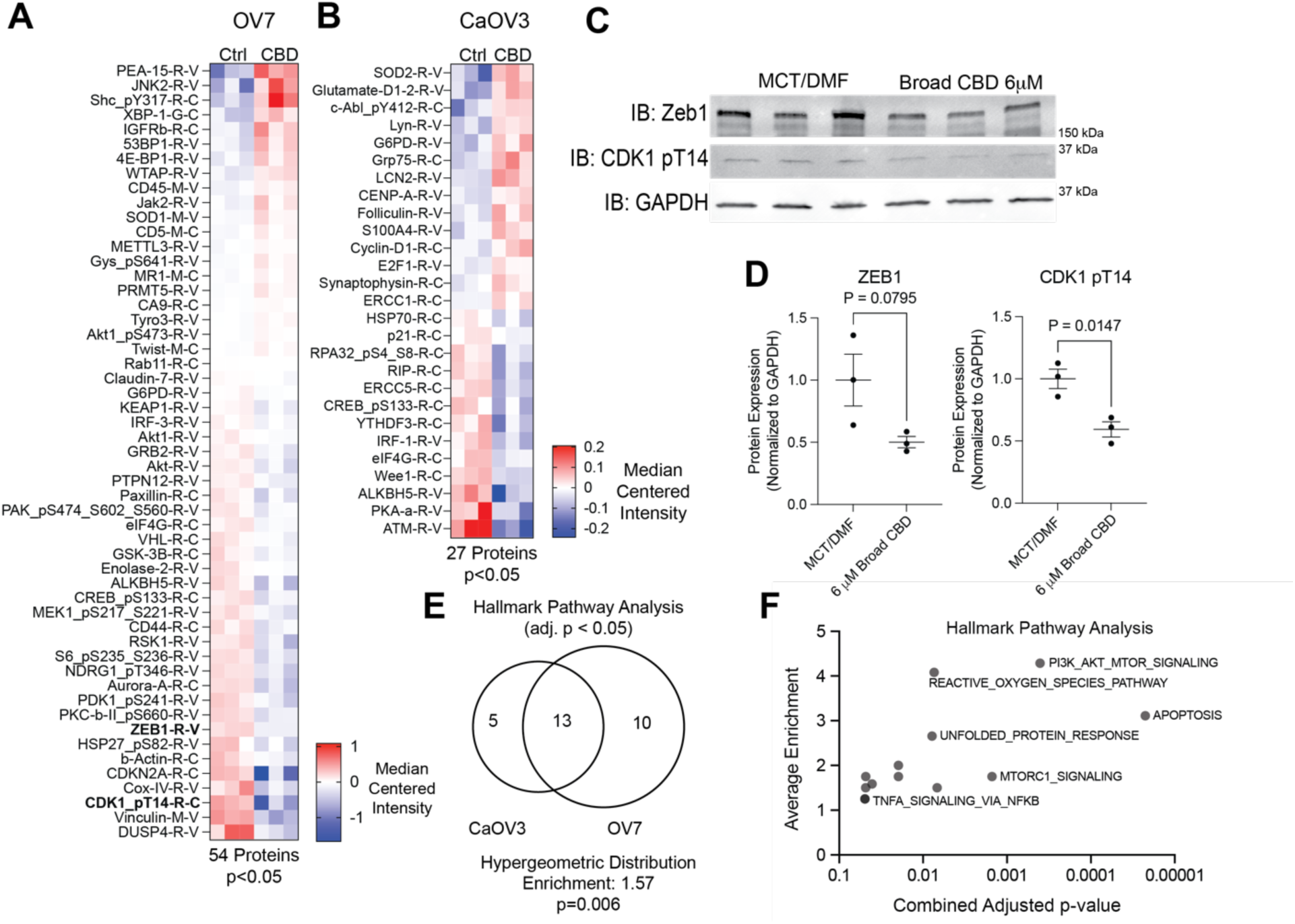
Defining CBD-induced protein signaling networks. A) OV7 and B) CaOV3 cells were treated with CBD for 72 hrs. Protein from treated cells was used for Reverse Phase Protein Array analysis. Heatmap of proteins with p<0.05. **C)** OV7 cells treated similarly as in A, were used for immunoblot against ZEB1 and CDK_pT14. Loading control, GAPDH. **D)** Densitometry of C. Signal intensity was normalized to GAPDH. **E)** Overlap of Hallmark Pathway enrichment analysis. **F)** Scatter plot of 13 overlapping pathways. Error bars, SD. Statistical test, unpaired t-test.

### Cell viability and apoptosis following treatment with CBD

Broad-spectrum CBD inhibits the growth of both OV7 and CaOV3 cells at higher concentrations (Figure 2), and the RPPA data indicated CBD-induced differential signaling (Figure 3). We next wanted to assess cell viability over time and measure cell death via cleaved caspase/7. Interestingly, our prior results found that full spectrum CBD had limited effect on 72 hr growth. Thus, we also wanted to determine if co-treatment with Full and Broad-spectrum CBD would overcome broad-spectrum alone-induced growth inhibition.

To more precisely understand cell dynamics following treatment with CBD, we used Incucyte live cell imaging. In a similar setup to the colony-forming viability assays, OV7 and CAOV3 cells were plated in 96 wells and treated with Broad, Full, and a 1:1 combination of full/broad spectrum CBD. We hypothesize that CBD-receptor interactions are driving the observed response, which led us to test whether treatment with full-spectrum may overcome or attenuate the effects of broad-spectrum. Consistently, in the OV7 and CaOV3, the broad spectrum CBD significantly reduced cell number at 30 μM concentrations. The treatment with the mixture of full:broad-spectrum CBD showed similar effects as broad alone. In both cell lines, the cell number dropped below the starting point (**Figure 4A-B**), suggesting CBD-induced cell death. Using a cleaved caspase 3/7 fluorescent dye, we next measure apoptosis over 72 hrs. In OV7, 6 and 30 μM broad-spectrum induced apoptosis within 12 hours, also when the peak apoptotic response to 30 μM broad-spectrum was observed (**Figure 4C-D**). In CaOV3, the CBD also induced an apoptotic response after 12 hours; however, the apoptosis peaked later at 48 hrs (**Figure 4E-F**). These data show that broad-spectrum CBD induces apoptosis, and this effect was not rescued by adding full-spectrum CBD.

**Figure 4.**
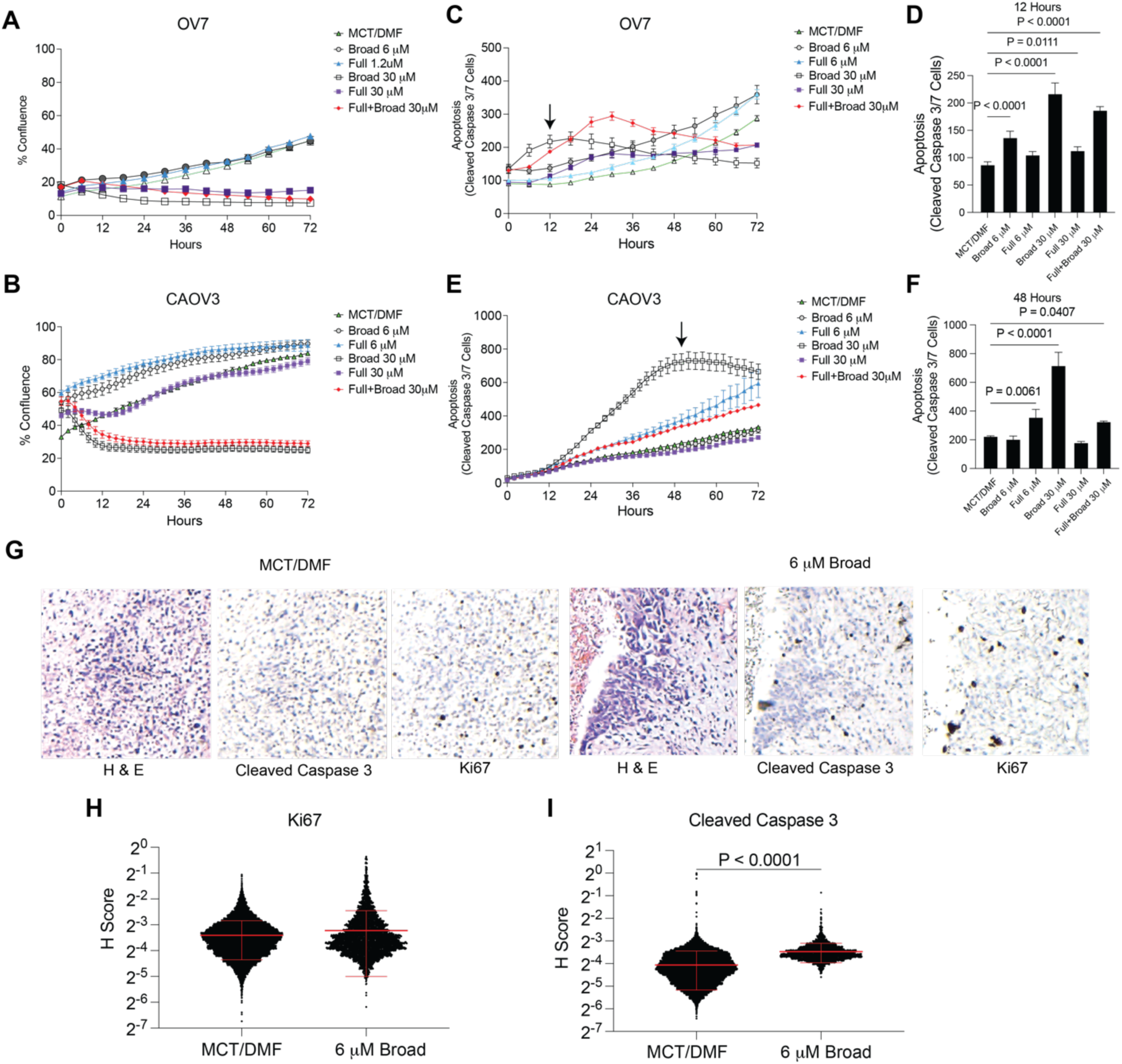
Broad spectrum CBD limits cell viability through the induction of apoptosis. **A)** Live imaging analysis of OV7 and **B)** CaOV3 cells following treatment with CBD. **C)** Evaluation of cleaved caspase 3/7 activity via fluorescence in OV7 cells over 72 hrs. **D)** Quantification of apoptotic OV7 cells at the 12 hr timepoint (arrow shown on C). **E)** Evaluation of cleaved caspase 3/7 activity via fluorescence in CaOV3 cells over 72 hrs. **F)** Quantification of apoptotic CaOV3 cells at the 48 hr timepoint (arrow shown on E). **G)** Ex vivo culture of a primary HGSC tumor. Representative images of H&E, Cleaved Caspase 3, and Ki67. **H)** Quantification of Ki67 histology score (H-score). **I)** Quantification of cleaved caspase 3 histology score (H-score). Error bars, SD. Statistical test, multicomparison ANOVA with Tukey correction (D, F) and unpaired t-test (H,I).

We next assessed the effect of broad-spectrum CBD on an *ex vivo* culture of a human primary HGSC tumor. The tumor was surgically resected and uniformly sliced. Tumor slices were cultured in vehicle control (DMF/MCT) or 6 μM Broad spectrum CBD for 72 hrs. Using immunohistochemistry (IHC), we measured proliferation and apoptosis via Ki67 and cleaved caspase 3, respectively. We scored the IHC using quantitative software. We observed no difference in Ki67 histology score (H-score); however, compared to vehicle treatment, there was a significant increase in cleaved caspase 3 signal (**Figure 4G-I**). These data demonstrate that in a primary tumor, CBD did not increase the growth of the tumor cells but induced cell death.

## DISCUSSION

The role of cannabinoids in oncology is still not fully understood, particularly in the realm of the effects cannabinoids such as CBD have on cell proliferation and treatment. There has been conflicting data regarding the possible beneficial and detrimental effects of cannabinoids on certain cancers, with a paucity of data in gynecologic malignancies. In this study, we used publicly available datasets to understand better CBD receptors’ expression and functional impacts in HGSC.

While CNR1/CNR2 is frequently discussed as the major receptors that modulate a downstream response for cannabinoids such as CBD, many other key receptors appear to play critical roles. When looking specifically at HGSC tumors and cell lines, FAAH, TRPV2, and TRPV4 showed the highest expression. TRPV2/TRPV4 are ion channels that are overexpressed in multiple tumor types and associated with tumor formation, progression, and metastasis. Overexpression of TRPV2/TRPV4 in lung cancer cell lines induced apoptosis via p38 MAPK^24^. In triple-negative breast cancers, longer relapse-free survival is associated with overexpression of TRPV4 ^25^. When looking at HGSC which had high expression compared to lower expression of the identified CBD receptors, the only one related to prognosis was low expression of FAAH. FAAH is a cannabinoid degradation enzyme that had a disease-specific survival benefit with an HR of 0.636 (p=0.0428). This was confirmed by evaluating CRISPR knockout in HGSC cells that, while not statistically significant, FAAH knockout demonstrated the lowest Chronos Dependency Score (Avg: −0.204, SD: 0.127), suggesting that loss of FAAH reduced cell viability. This idea of increasing cannabinoids in the system by inhibiting FAAH is being explored in both neurological diseases and in the realm of oncology ^26,27^.

Treatment of HGSC cells with broad-spectrum CBD showed decreased cell viability by inducing PI3K/AKT/MTOR apoptotic pathways. The broad-spectrum CBD, a purer formulation compared to full-spectrum CBD, which contains multiple cannabinoids, exhibited greater antitumor properties. It is suspected that while CBD may have antitumor properties, perhaps other cannabinoids that were in the full spectrum of CBD may have an antagonistic effect with cisplatin. More studies will be needed to define individual cannabinoids and their roles within oncology. Significant information regarding the mechanism of action that contributed to CBD decreasing proliferation and cell death was gained in this study. RPPA analysis identified significant modulation of proteins associated with the PI3K/AKT/mTOR pathways, which are involved in cell proliferation and survival. Dysregulation of this pathway is a hallmark of carcinogenesis and is a target of many cancer therapies that aim to inhibit its activity^28^. While both cancer cell lines that were used in this study showed different proteins that were modulated by CBD treatment, overall downstream effects were similar, implying that it is not one signaling mechanism involved with CBD but a broader impact on cellular stress and apoptosis.

The antitumor properties of CBD were noted in a dose-dependent fashion and only appreciated at higher concentrations. While further studies are needed to determine the maximum safe concentrations for human CBD consumption, the dosing required to achieve significant antitumor effects in this study was generally lower than typical oral dosing recommendations^29^. However, intravenous formulations that allow for better absorption may allow for closer dosing and have been well tolerated in patients being treated for seizures^30^. In this study, cells treated with CBD did not significantly alter the IC50 concentrations of cisplatin and paclitaxel. This finding addresses a common concern among patients taking oral CBD supplements, suggesting that CBD is unlikely to meaningfully impact the efficacy of their primary cancer treatment. However, further studies are needed to definitively confirm this.

This study contributes to the growing research on cannabinoids in oncology, specifically focusing on CBD in HGSC. While the insights gained from in vitro cell line and *ex vivo* tumor experiments are promising, further exploration is needed. In vivo studies will be essential to confirm CBD’s dosing, efficacy, and safety as a potential therapeutic agent in HGSC.

## ACKNOWLEDGEMENTS

We acknowledge philanthropic contributions from D. Thomas and Kay L. Dunton Endowed Chair in Ovarian Cancer Research, the McClintock-Addlesperger Family, Karen M. Jennison, Don and Arlene Mohler Johnson Family, Michael Intagliata, Duane and Denise Suess, Mary Normandin, and Donald Engelstad. The study was funded entirely through the University of Colorado Department of OB/GYN Academic Enrichment Fund (AG). This study utilized the University of Colorado Cancer Center shared resources, partly supported by the National Cancer Institute through Cancer Center Support Grant P30CA046934.

