## Supplemental Table 1 for "Cannabidiol promotes apoptosis and downregulation of oncogenic factors"

**Supplemental Table 1. CBD-dependent signaling**. CaOV3 and OV7 treated with MCT/DMF or 6 mM Broad spectrum CBD for 24 hrs. Protein collected for reverse phase protein array analysis. Values = median centered by antibodies.

**Supplemental Table 1, Greenwood et al 2024**

| | CaOV3 | | | | | |
|---|---|---|---|---|---|---|
| **Antibody Name** | **DMF/MCT** | **DMF/MCT** | **DMF/MCT** | **6 uM Broad Spectrum CBD** | **6 uM Broad Spectrum CBD** | **6 uM Broad Spectrum CBD** |
| Wee1 | 0.0374 | 0.0550 | 0.0707 | -0.0427 | -0.0564 | -0.0517 |
| ATM | 0.0901 | 0.1955 | 0.1922 | -0.1425 | -0.1061 | -0.2058 |
| LCN2 | -0.0345 | -0.0393 | -0.0499 | 0.1005 | 0.0803 | 0.0310 |
| Lyn | -0.0706 | -0.0277 | -0.0905 | 0.0372 | 0.0236 | 0.0375 |
| CENP-A | -0.0197 | -0.0437 | -0.0505 | 0.0144 | 0.0220 | 0.0460 |
| ERCC5 | 0.0163 | 0.0453 | 0.0519 | -0.1017 | -0.0323 | -0.0688 |
| Glutamate-D1-2 | -0.0332 | -0.1162 | -0.1032 | 0.0278 | 0.0883 | 0.0619 |
| Grp75 | -0.0659 | -0.0429 | -0.0255 | 0.0686 | 0.1153 | 0.0348 |
| RIP | 0.0509 | 0.0234 | 0.0360 | -0.1165 | -0.0275 | -0.0875 |
| IRF-1 | 0.0106 | 0.0441 | 0.0754 | -0.0804 | -0.0266 | -0.0994 |
| G6PD | -0.0626 | -0.0895 | -0.0175 | 0.0581 | 0.0143 | 0.0885 |
| E2F1 | -0.0129 | -0.0348 | -0.0242 | 0.0396 | 0.0092 | 0.0094 |
| SOD2 | -0.0383 | -0.0879 | -0.1889 | 0.0605 | 0.0892 | 0.0348 |
| ERCC1 | -0.0190 | -0.0175 | -0.0181 | 0.0486 | 0.0030 | 0.0412 |
| Folliculin | -0.0383 | -0.0132 | -0.0462 | 0.0864 | 0.0091 | 0.0618 |
| eIF4G | 0.0048 | 0.0641 | 0.0618 | -0.0547 | -0.0208 | -0.0167 |
| S100A4 | -0.0686 | -0.0122 | -0.0152 | 0.0763 | 0.0429 | 0.0206 |
| ALKBH5 | 0.0615 | 0.1172 | 0.0572 | -0.2426 | -0.0683 | -0.0480 |
| c-Abl_pY412 | -0.1489 | -0.0604 | -0.0221 | 0.0294 | 0.0445 | 0.0406 |
| Cyclin-D1 | -0.0140 | -0.0149 | -0.0486 | 0.0087 | 0.0541 | 0.0830 |
| RPA32_pS4_S8 | 0.0556 | 0.0152 | 0.0245 | -0.1171 | -0.0193 | -0.0436 |
| CREB_pS133 | 0.0642 | 0.0490 | 0.0013 | -0.1047 | -0.0045 | -0.0983 |
| PKA-a | 0.0506 | 0.0422 | 0.2055 | -0.0458 | -0.0463 | -0.1465 |
| Synaptophysin | -0.0416 | -0.0200 | -0.0017 | 0.0654 | 0.0280 | 0.0109 |
| HSP70 | 0.0136 | 0.0270 | 0.0026 | -0.0615 | -0.0058 | -0.0706 |
| p21 | -0.0020 | 0.0277 | 0.0302 | -0.0034 | -0.0214 | -0.0187 |
| YTHDF3 | 0.0349 | 0.0098 | 0.0785 | -0.1461 | -0.0139 | -0.0770 |
| ATM_pS1981 | -0.0122 | -0.1105 | -0.0992 | 0.0333 | -0.0038 | 0.0909 |
| RBM15 | 0.0592 | 0.0687 | 0.0053 | -0.1295 | -0.0085 | -0.0572 |
| MEK1 | -0.0037 | 0.0320 | 0.0230 | -0.1281 | -0.0123 | -0.1310 |
| FANCD2 | -0.0619 | -0.0080 | -0.1065 | 0.0145 | 0.0128 | 0.0836 |
| FASN | -0.0039 | 0.0516 | 0.1190 | -0.0647 | -0.0780 | 0.0005 |
| Rb_pS807_S811 | 0.0741 | 0.0080 | 0.1653 | -0.0963 | -0.0122 | -0.0216 |
| PAR | -0.0216 | 0.3579 | 0.3073 | -0.1611 | -0.3409 | 0.0181 |
| Notch1 | -0.0030 | -0.0310 | -0.0379 | -0.0024 | 0.0621 | 0.0219 |
| SOD1 | 0.0852 | 0.0015 | 0.1799 | -0.0686 | -0.0057 | -0.1927 |
| Stat5a | 0.0519 | -0.0031 | 0.0773 | -0.0425 | -0.0011 | -0.0177 |
| Myt1 | 0.0019 | 0.0077 | 0.0720 | -0.1037 | -0.1364 | 0.0007 |

| | | | | | | |
|---|---|---|---|---|---|---|
| eIF4E_pS209 | 0.0242 | -0.0054 | 0.0142 | -0.0630 | 0.0013 | -0.0483 |
| PKC-a-b-II_pT638_T641 | -0.0135 | 0.4325 | 0.4531 | -0.1610 | -0.0024 | -0.0174 |
| ATR_pS428 | 0.0321 | 0.0058 | 0.0025 | -0.0651 | -0.0770 | 0.0068 |
| Sox17 | -0.0262 | -0.0004 | -0.0244 | 0.0130 | -0.0037 | 0.0462 |
| TSC1 | -0.0141 | 0.0018 | -0.0560 | -0.0088 | 0.0617 | 0.0740 |
| BRD4 | 0.0429 | -0.0149 | 0.0548 | -0.1445 | 0.0108 | -0.1932 |
| DJ1 | -0.0110 | 0.1337 | 0.1462 | -0.0668 | -0.0050 | -0.0023 |
| Akt2 | 0.0357 | 0.0042 | 0.0174 | -0.0555 | -0.0083 | -0.0007 |
| CD44 | -0.0375 | 0.0200 | -0.1701 | -0.0135 | 0.1370 | 0.1657 |
| YAP | -0.0240 | 0.1196 | 0.1330 | -0.0630 | -0.0942 | 0.0205 |
| c-Jun_pS73 | -0.0153 | 0.0030 | -0.0221 | -0.0067 | 0.0238 | 0.0276 |
| Erk5 | -0.1607 | -0.0002 | 0.0006 | 0.2021 | -0.0079 | 0.1364 |
| IRS1 | -0.0514 | 0.5617 | 0.3207 | -0.1677 | 0.0354 | -0.1730 |
| EphA2_pS897 | -0.2796 | 0.0058 | -0.0428 | 0.0007 | 0.1097 | 0.1816 |
| CIITA | -0.0352 | -0.0240 | 0.0044 | 0.0507 | 0.0035 | 0.0049 |
| PHGDH | -0.0345 | -0.0810 | 0.0115 | 0.0520 | 0.0272 | -0.0023 |
| Src_pY527 | -0.0264 | 0.1215 | 0.2110 | 0.0210 | -0.0549 | -0.1124 |
| ZEB1 | -0.0495 | 0.0183 | -0.0863 | 0.1143 | 0.0473 | -0.0100 |
| Cyclin-D3 | 0.0583 | -0.0058 | 0.0819 | -0.2059 | 0.0017 | -0.0481 |
| MSI2 | -0.0259 | 0.0713 | 0.0831 | -0.0706 | 0.0099 | -0.0502 |
| Cdc6 | -0.0050 | -0.0128 | -0.0090 | 0.0612 | -0.0110 | 0.0520 |
| IDO | -0.0414 | -0.0343 | 0.0129 | 0.0161 | 0.0496 | -0.0008 |
| EphA2_pY588 | -0.0948 | -0.0134 | -0.0038 | 0.0112 | 0.0211 | 0.0172 |
| NF-kB-p65_pS536 | 0.0528 | -0.0298 | 0.1398 | -0.1206 | 0.0256 | -0.2160 |
| ATR | -0.0119 | -0.0004 | -0.0458 | 0.0040 | 0.0136 | 0.0858 |
| P-Cadherin | -0.0115 | 0.0384 | 0.0440 | -0.0436 | -0.0211 | 0.0080 |
| p70-S6K_pT389 | -0.0249 | 0.2885 | 0.2286 | 0.0195 | -0.0663 | -0.0078 |
| Wee1_pS642 | -0.0001 | -0.0295 | -0.0113 | 0.0335 | 0.0361 | -0.0123 |
| Akt | -0.0073 | 0.0419 | 0.0879 | -0.0733 | -0.0087 | -0.3247 |
| Hexokinase-II | -0.1357 | 0.5898 | 0.6017 | -0.5689 | 0.0804 | -0.0757 |
| Melanoma-gp100 | -0.0721 | 0.0039 | -0.0231 | 0.0846 | -0.0080 | 0.0155 |
| CtIP | -0.1368 | 0.0055 | -0.0473 | 0.0437 | -0.0090 | 0.0137 |
| IGFBP2 | -0.1857 | 0.0353 | -0.1825 | 0.0299 | 0.0338 | -0.0146 |
| Stat1_pY701 | -0.0380 | 0.0040 | -0.0035 | 0.0608 | 0.0158 | -0.0006 |
| CD20 | -0.0854 | 0.0358 | -0.3266 | 0.1124 | -0.0399 | 0.1800 |
| TRIM24 | 0.0212 | -0.0522 | -0.0412 | 0.0149 | 0.0462 | 0.0004 |
| CA9 | 0.0485 | 0.0485 | -0.0106 | -0.0386 | 0.0045 | 0.0002 |
| ERRalpha | 0.1057 | -0.0073 | 0.0253 | -0.0103 | 0.0032 | -0.0824 |
| PGM1 | 0.0873 | -0.0415 | 0.1427 | -0.0970 | 0.0374 | -0.1117 |
| CD31 | -0.0356 | -0.0034 | 0.0038 | 0.0537 | 0.1597 | -0.0036 |
| UBAC1 | 0.0535 | -0.1476 | -0.3024 | 0.3017 | -0.0110 | 0.0158 |
| BRCA1 | -0.0800 | 0.0233 | -0.0229 | 0.1152 | 0.0844 | -0.0250 |
| DDR1_pY513 | -0.0706 | 0.0198 | -0.0516 | 0.0438 | 0.0060 | -0.0013 |
| PIP4K2B | -0.0079 | -0.0044 | -0.0208 | 0.0304 | -0.0159 | 0.0523 |
| Folate-Binding-Protein | 0.0304 | 0.1768 | -0.0050 | -0.0285 | 0.0018 | -0.0442 |
| TFRC | -0.0796 | 0.2207 | 0.3134 | -0.1935 | -0.0267 | 0.0314 |
| PDK1_pS241 | -0.0316 | 0.0193 | 0.0935 | -0.1509 | -0.3336 | 0.0380 |
| JAB1 | 0.1279 | 0.0737 | -0.0151 | -0.0116 | 0.0118 | -0.0098 |
| Annexin-VII | 0.0370 | -0.0232 | 0.0733 | -0.0415 | 0.0191 | -0.1328 |
| c-Kit | 0.0918 | -0.0378 | 0.3810 | -0.0096 | 0.0124 | -0.3405 |
| CD4 | -0.1136 | 0.0198 | -0.0307 | 0.0846 | 0.0226 | -0.0115 |

| | | | | | | |
|---|---|---|---|---|---|---|
| DM-K9-Histone-H3 | 0.0403 | 0.0248 | -0.0082 | -0.0292 | 0.0030 | 0.0017 |
| Rab11FIP1 | 0.0003 | -0.0624 | -0.0327 | 0.2823 | -0.0163 | 0.0578 |
| MSH6 | 0.0118 | -0.0241 | 0.1461 | -0.1735 | 0.0323 | -0.0830 |
| PTPN12 | 0.0204 | 0.0973 | -0.0071 | 0.0059 | -0.0031 | -0.0678 |
| Myosin-IIa | -0.0225 | 0.0824 | 0.0180 | -0.1749 | 0.0065 | -0.0363 |
| LRP6_pS1490 | -0.0150 | 0.0212 | 0.0967 | -0.0554 | -0.0250 | 0.0116 |
| PDK1 | 0.0376 | -0.0113 | 0.0547 | -0.0983 | 0.0071 | -0.0027 |
| PAK_pS474_S602_S560 | -0.0174 | 0.0124 | 0.0564 | 0.0120 | -0.0356 | -0.1068 |
| c-Met_pY1234_Y1235 | 0.0044 | -0.0177 | -0.0634 | 0.0120 | -0.0092 | 0.0164 |
| MYH11 | 0.1657 | 0.0779 | -0.0342 | -0.1026 | 0.0301 | -0.0254 |
| ARID1A | -0.0028 | 0.0048 | -0.0044 | -0.0289 | 0.0026 | -0.0185 |
| H2AX_pS139 | 0.0694 | -0.0168 | 0.0381 | -0.0247 | 0.0126 | -0.1895 |
| Rictor | -0.0324 | 0.0790 | 0.1223 | 0.0027 | 0.0002 | -0.0444 |
| UVRAG | 0.0120 | -0.0011 | 0.0016 | -0.0636 | -0.0422 | 0.0151 |
| p27-Kip1 | -0.0104 | 0.0129 | -0.0010 | -0.0667 | 0.0147 | -0.0671 |
| TEAD | -0.0233 | 0.0204 | 0.0130 | -0.0868 | -0.0812 | 0.0198 |
| A-Raf | 0.0439 | -0.0528 | -0.2041 | 0.0030 | -0.0001 | 0.2112 |
| Cdc42 | 0.0301 | -0.0071 | 0.0212 | -0.0621 | 0.0030 | 0.0010 |
| NDRG1_pT346 | -0.1230 | 0.5477 | 0.4025 | -0.0483 | 0.0432 | -0.0385 |
| B-Raf | 0.0046 | -0.0013 | 0.0343 | -0.0317 | -0.1560 | 0.0097 |
| SIRP-alpha | -0.0723 | 0.1422 | 0.2161 | -0.1248 | 0.0563 | -0.0730 |
| ATRX | 0.0439 | -0.0377 | 0.3660 | -0.0497 | 0.0335 | -0.2053 |
| A-Raf_pS299 | 0.0128 | -0.0251 | -0.0669 | 0.0719 | 0.0547 | -0.0359 |
| MEK1_pS217_S221 | -0.3561 | -0.0326 | 0.0330 | -0.0641 | 0.0750 | 0.2878 |
| E-Cadherin | -0.0091 | 0.0869 | 0.4999 | 0.0037 | -0.0071 | -0.0731 |
| SGK1 | -0.0754 | 0.0270 | -0.0462 | 0.0302 | -0.0274 | 0.1673 |
| SDHA | 0.0467 | -0.1002 | -0.0847 | 0.0081 | 0.0401 | 0.0072 |
| GCN5L2 | -0.0123 | -0.3464 | 0.0009 | 0.1792 | 0.1001 | -0.0862 |
| p44-42-MAPK | -0.0136 | 0.0012 | 0.0766 | -0.1150 | -0.0189 | 0.0107 |
| MIG6 | -0.1034 | -0.3094 | 0.0897 | 0.2591 | 0.0669 | -0.0622 |
| RSK | -0.0286 | 0.1002 | 0.0410 | -0.0677 | -0.0120 | 0.0167 |
| PLK1 | 0.0079 | -0.0119 | -0.1256 | 0.0183 | 0.1348 | -0.0307 |
| KEAP1 | -0.0175 | 0.0052 | 0.0397 | -0.0961 | -0.1222 | 0.0393 |
| MLKL | -0.0428 | 0.1275 | 0.1562 | 0.0208 | -0.0179 | -0.0088 |
| ACVRL1 | 0.0142 | -0.0147 | -0.0380 | -0.0043 | 0.0105 | 0.0196 |
| WIPI2 | -0.0240 | 0.0386 | 0.1825 | 0.0186 | -0.0172 | -0.0541 |
| Caspase-8-cleaved | 0.0029 | -0.0067 | -0.0593 | 0.0114 | -0.0084 | 0.0131 |
| Akt_pS473 | -0.1016 | 0.0748 | -0.1315 | -0.0307 | 0.1316 | 0.0460 |
| DUSP6 | -0.0747 | 0.1229 | 0.1128 | -0.0752 | 0.0587 | -0.1984 |
| HSP27 | 0.0565 | 0.0760 | -0.0303 | 0.0377 | -0.0619 | -0.0446 |
| HER2_pY1248 | -0.0880 | 0.0137 | -0.0132 | -0.0208 | 0.0283 | 0.0378 |
| LAD1 | 0.1280 | 0.0275 | -0.0271 | 0.0468 | -0.1038 | -0.0516 |
| c-Abl | 0.0013 | -0.0136 | 0.1744 | 0.0521 | -0.0956 | -0.0759 |
| TAZ | 0.1533 | 0.0238 | -0.0234 | -0.1240 | 0.0452 | -0.0325 |
| Atg3 | 0.0658 | -0.0410 | 0.1525 | 0.0347 | -0.0319 | -0.0500 |
| COG3 | -0.0146 | 0.0210 | 0.0382 | -0.0259 | -0.0093 | 0.0111 |
| PD-1 | -0.0368 | 0.0301 | -0.0457 | 0.0123 | -0.0095 | 0.0579 |
| AMPKa_pT172 | 0.1062 | -0.1463 | -0.4150 | -0.0552 | 0.1048 | 0.0705 |
| PTEN | 0.0133 | -0.0108 | 0.2186 | -0.0374 | 0.0066 | -0.0152 |
| Gli1 | -0.0053 | -0.0070 | 0.0837 | -0.0485 | -0.0345 | 0.0214 |
| Bcl2 | -0.4872 | 0.0744 | -0.1057 | -0.0679 | 0.0886 | 0.0880 |

| | | | | | | |
|---|---|---|---|---|---|---|
| TRIP13 | -0.0420 | -0.1039 | 0.0306 | 0.1921 | -0.0608 | 0.0509 |
| BAP1 | 0.0652 | -0.0094 | 0.0098 | -0.0114 | 0.0207 | -0.1720 |
| 14-3-3-beta | -0.0905 | -0.0102 | 0.0106 | 0.0892 | -0.0663 | 0.1205 |
| BiP-GRP78 | 0.0314 | 0.0534 | -0.0196 | 0.0270 | -0.0276 | -0.0408 |
| CSK | 0.0000 | -0.0124 | -0.1414 | -0.0658 | 0.0432 | 0.1091 |
| MCT4 | 0.2573 | -0.2696 | 0.4823 | -0.5827 | 0.4348 | -1.3220 |
| RPA32 | 0.1018 | -0.0600 | 0.6473 | -0.0625 | 0.0121 | -0.0074 |
| ER-a_pS118 | 0.0267 | -0.0434 | -0.0390 | -0.0074 | 0.0128 | 0.0227 |
| Axl | -0.0548 | -0.1338 | 0.0433 | 0.8798 | 0.0416 | -0.0400 |
| Jak2 | 0.0433 | -0.0405 | -0.1196 | -0.0243 | 0.0472 | 0.0396 |
| GSK-3a-b | 0.0415 | -0.0132 | 0.0355 | 0.0064 | -0.0173 | 0.0090 |
| eEF2 | 0.0115 | -0.0238 | 0.0514 | -0.6283 | -0.5978 | 0.2635 |
| Claudin-7 | 0.1071 | -0.1194 | 0.2908 | -0.1793 | 0.1725 | -0.4567 |
| PAI-1 | 0.0415 | -0.0221 | -0.1042 | 0.5444 | -0.0136 | 0.0183 |
| PAK4 | 0.0580 | -0.0507 | -0.1309 | 0.0097 | 0.0543 | 0.0056 |
| Tuberin | 0.0509 | -0.0287 | 0.0513 | -0.0356 | 0.0182 | -0.0135 |
| DRP1 | -0.3034 | -0.0703 | 0.0666 | 0.0664 | -0.0335 | 0.0382 |
| Histone-H3_pS10 | -0.0547 | 0.0465 | -0.0468 | -0.0036 | 0.0064 | 0.0907 |
| Coup-TFII | -0.1056 | -0.0157 | 0.0173 | 0.0176 | -0.0148 | 0.0227 |
| GCLM | -0.0276 | 0.0153 | -0.0536 | 0.0533 | 0.0164 | -0.0301 |
| Smad4 | -0.0072 | -0.0032 | 0.0029 | -0.0003 | -0.0009 | 0.0064 |
| DAPK2 | -0.0134 | 0.0273 | 0.0603 | -0.0025 | -0.0029 | 0.0076 |
| Atg5 | -0.0542 | 0.0495 | 0.3388 | 0.0000 | 0.0028 | -0.0675 |
| Fibronectin | -0.1040 | 0.0278 | -0.0274 | 0.1860 | 0.0468 | -0.0651 |
| Mitofusin-1 | -0.0185 | 0.0062 | 0.1246 | -0.0630 | -0.1146 | 0.0630 |
| Cyclophilin-F | 0.0544 | -0.0667 | -0.0995 | -0.1167 | 0.2826 | 0.1337 |
| IGFBP3 | 0.1190 | 0.0716 | -0.0665 | -0.1215 | 0.0455 | -0.0408 |
| NRF2 | -0.0197 | 0.0074 | -0.1643 | 0.1232 | 0.0073 | -0.0611 |
| IGF1R_pY1135_Y1136 | 0.0376 | 0.0097 | -0.0093 | -0.0745 | 0.0181 | -0.0058 |
| STING | 0.0428 | -0.0551 | 0.0800 | -0.3614 | 0.1061 | -0.1290 |
| FAK_pY397 | -0.2033 | 0.0812 | -0.0912 | -0.0273 | 0.0474 | 0.0426 |
| MelanA | -0.0354 | -0.0060 | 0.0065 | 0.0159 | -0.0247 | 0.0708 |
| Heregulin | 0.0655 | -0.0500 | -0.2292 | -0.0573 | 0.0950 | 0.0584 |
| C-Raf | 0.0447 | -0.0091 | 0.0015 | -0.2314 | -0.0047 | 0.0168 |
| PR | -0.0355 | -0.0016 | 0.0131 | 0.0361 | -0.0025 | 0.0020 |
| TIGAR | -0.0359 | 0.0236 | -0.0286 | -0.0294 | 0.0225 | 0.0914 |
| b-Actin | 0.0665 | -0.0261 | 0.0092 | -0.0018 | 0.0096 | -0.0835 |
| Chk2_pT68 | 0.0455 | -0.0354 | -0.0477 | 0.0990 | 0.0225 | -0.0178 |
| SFRP1 | 0.0649 | -0.0019 | 0.0023 | -0.0680 | 0.0240 | 0.0008 |
| IL-6 | -0.1165 | -0.0117 | 0.0430 | 0.0725 | -0.0066 | 0.0113 |
| cGAS | 0.0920 | 0.0738 | -0.0565 | -0.0491 | -0.0001 | 0.0048 |
| Mnk1 | 0.1074 | 0.1087 | -0.1012 | -0.1758 | 0.0273 | -0.0226 |
| Smad3 | 0.0564 | -0.1921 | -0.0260 | 0.0697 | 0.0107 | -0.0060 |
| Bid | -0.0130 | 0.0070 | -0.0379 | -0.0224 | 0.0305 | 0.0095 |
| PRMT5 | 0.0089 | -0.0035 | -0.0387 | 0.0100 | -0.0224 | 0.0888 |
| 14-3-3-zeta | -0.7158 | 0.0197 | -0.0075 | -0.0578 | 0.0756 | 0.0167 |
| IRS2 | 0.0809 | -0.0245 | -0.0788 | 0.0310 | 0.2721 | -0.0189 |
| Rad17_pS645 | 0.0594 | -0.0336 | -0.0691 | -0.0086 | 0.0114 | 0.1219 |
| Mcl-1 | 0.0987 | -0.1299 | -0.1185 | -0.0795 | 0.1330 | 0.0948 |
| MMP14 | -0.3129 | 0.6998 | 0.3700 | 0.2779 | -0.3042 | -0.2626 |
| Notch1-cleaved | 0.0062 | 0.0036 | -0.0032 | -0.0247 | 0.0031 | 0.0007 |

| | | | | | | |
|---|---|---|---|---|---|---|
| Twist | 0.0188 | -0.0121 | -0.0176 | 0.0033 | 0.0079 | 0.0121 |
| EphA2 | 0.1283 | 0.0176 | -0.0172 | -0.1111 | -0.5584 | 0.1719 |
| Stat3 | 0.0135 | -0.0258 | 0.1002 | -0.2179 | 0.1156 | -0.1133 |
| C-Raf_pS338 | -0.0154 | 0.0031 | 0.0067 | 0.1279 | -0.2444 | -0.2503 |
| FGFR2 | -0.1109 | 0.0057 | -0.0053 | 0.0721 | -0.0673 | 0.0479 |
| Atg4B | -0.0224 | 0.0101 | 0.0135 | -0.0697 | -0.0491 | 0.0286 |
| PEA-15 | 0.0405 | -0.0848 | 0.1219 | 0.0190 | -0.1967 | -0.0037 |
| PAX8 | 0.0190 | -0.0179 | -0.0119 | -0.0073 | 0.0087 | 0.0360 |
| DNMT1 | 0.0735 | -0.0027 | 0.0031 | -0.0087 | -0.3579 | 0.0654 |
| FOXM1 | 0.0102 | -0.0225 | -0.0501 | 0.0300 | -0.0506 | 0.0690 |
| CD49b | 0.0025 | 0.0117 | -0.3919 | -0.0052 | 0.0489 | -0.0402 |
| GGPS1 | 0.0133 | -0.0065 | 0.0070 | -0.0154 | 0.0080 | -0.0039 |
| GAPDH | 0.0457 | -0.2119 | 0.1104 | -1.4668 | -0.0617 | 0.0697 |
| MAPK_pT202_Y204 | 0.1041 | -0.0548 | -0.0476 | -0.0612 | 0.0444 | 0.6946 |
| SGK3 | -0.1802 | 0.0067 | 0.0028 | 0.0973 | -0.0612 | 0.0065 |
| NQO1 | 0.0070 | -0.0194 | -0.1877 | -0.1440 | 0.0834 | 0.1665 |
| PERK | -0.0174 | -0.0072 | 0.0060 | 0.0401 | 0.0253 | -0.0260 |
| Calnexin | -0.2409 | 0.0088 | -0.0084 | 0.1058 | 0.0573 | -0.1161 |
| FABP5 | 0.0242 | -0.0138 | -0.0031 | 0.1548 | -0.0002 | -0.0024 |
| Rab25 | -0.0164 | 0.0041 | 0.0798 | -0.0253 | 0.0202 | -0.0154 |
| Syk | 0.6824 | -0.0322 | 0.0326 | -0.3882 | 0.2995 | -0.0455 |
| DDB-1 | 0.0758 | -0.0452 | 0.0483 | -0.0284 | 0.0011 | 0.0036 |
| SCD | -0.0315 | 0.1360 | -0.1951 | 0.0262 | -0.0568 | 0.3877 |
| SHP-2_pY542 | 0.0425 | 0.0417 | -0.0592 | -0.1305 | 0.0173 | -0.0126 |
| Cyclin-E1 | 0.0964 | -0.0842 | -0.1380 | 0.0755 | 0.0339 | -0.0291 |
| VHL | -0.0549 | -0.0192 | 0.0407 | 0.0600 | 0.0151 | -0.0143 |
| CRABP2 | 0.0302 | 0.0411 | -0.0295 | 0.0231 | -0.0203 | -0.0360 |
| CD68 | 0.0123 | 0.0223 | -0.0195 | -0.0462 | 0.0120 | -0.0073 |
| WTAP | 0.0260 | -0.1035 | 0.2129 | 0.0302 | -0.0273 | -0.1491 |
| Enolase-2 | 0.0109 | -0.0411 | 0.0231 | 0.0027 | 0.0001 | -0.1719 |
| GSK-3a-b_pS21_S9 | 0.1149 | 0.0539 | -0.0638 | -0.0458 | -0.0072 | 0.0119 |
| DUSP4 | 0.0324 | -0.0146 | -0.0465 | -0.0162 | 0.0391 | 0.0230 |
| GCLC | 0.0386 | -0.2371 | 0.4559 | -0.0440 | 0.1018 | -0.5373 |
| XPA | 0.0240 | 0.0001 | -0.0719 | -0.0346 | 0.1233 | 0.0083 |
| p38-MAPK | -0.1142 | 0.1019 | 0.2455 | 0.1850 | -0.1403 | -0.2459 |
| Raptor | -0.0006 | -0.0117 | 0.0405 | -0.0351 | 0.0200 | -0.0156 |
| GRB2 | -0.0372 | 0.0560 | 0.0543 | -0.0104 | 0.0133 | -0.0120 |
| Stat3_pY705 | -0.0694 | 0.0369 | -0.0183 | -0.0152 | 0.0151 | 0.0378 |
| Tyro3 | 0.0087 | 0.1016 | -0.0375 | -0.0621 | -0.0247 | 0.0325 |
| PAX6 | 0.0301 | -0.0044 | -0.1775 | 0.0245 | 0.0003 | -0.0093 |
| Lck | 0.0138 | -0.0709 | 0.2061 | 0.0269 | -0.0952 | -0.0115 |
| HER3 | 0.2189 | 0.2208 | -0.2204 | 0.2356 | -0.2693 | -0.3249 |
| HNRNPK | -0.0155 | 0.0365 | 0.0041 | 0.0298 | -0.0205 | -0.0587 |
| GSK-3B | 0.0008 | -0.0131 | 0.0566 | -0.0613 | 0.0613 | -0.0703 |
| Smad1 | 0.0393 | -0.0090 | -0.1287 | 0.0155 | 0.0423 | -0.0273 |
| PLC-gamma1_pS1248 | 0.0386 | -0.0048 | -0.0218 | 0.0113 | 0.0645 | 0.0001 |
| MSH2 | -0.0346 | -0.0001 | 0.1075 | 0.0066 | -0.0720 | 0.0164 |
| GRB7 | -0.0085 | -0.0038 | -0.0198 | 0.0169 | 0.0351 | -0.0339 |
| CD29 | 0.0382 | 0.0125 | -0.0277 | -0.0477 | 0.0135 | -0.0042 |
| PARG | -0.0449 | 0.0596 | -0.0646 | 0.0396 | 0.0484 | -0.0311 |
| 4E-BP1_pT37_T46 | 0.0715 | -0.0850 | 0.0953 | 0.0564 | -0.1017 | -0.0411 |

| | | | | | | |
|---|---|---|---|---|---|---|
| MRAP | -0.0084 | 0.0122 | -0.0030 | -0.0549 | -0.0226 | 0.0239 |
| Ezh2 | 0.0415 | 0.0062 | -0.0304 | 0.0003 | -0.2545 | 0.0534 |
| MDM2_pS166 | 0.0090 | -0.0013 | 0.0017 | 0.0094 | -0.0459 | 0.0055 |
| Oct-4 | 0.0672 | -0.0014 | -0.0114 | -0.0775 | -0.0027 | 0.0402 |
| RIP3 | -0.0565 | 0.0706 | 0.0335 | -0.0651 | -0.0340 | 0.0387 |
| Gys_pS641 | 0.0361 | -0.0061 | 0.0066 | -0.0371 | -0.0131 | 0.0337 |
| LC3A-B | -0.1380 | -0.2222 | 0.1809 | 0.1326 | -0.1648 | 0.2219 |
| MMP2 | -0.0180 | 0.0148 | -0.0264 | 0.0127 | -0.0223 | 0.0190 |
| DDR1 | 0.0408 | -0.0119 | -0.0358 | 0.0184 | 0.1120 | -0.6800 |
| PI3K-p85 | 0.0854 | 0.0068 | -0.0730 | -0.1771 | -0.0110 | 0.0368 |
| FTO | 0.0133 | -0.0020 | -0.2897 | -0.0866 | 0.0173 | 0.0103 |
| Complex-II-Subunit | 0.0294 | -0.0199 | 0.0203 | -0.0337 | 0.0294 | -0.0164 |
| EVI1 | -0.0205 | 0.0518 | -0.0708 | 0.0151 | 0.0297 | -0.0071 |
| Vinculin | 0.0593 | -0.0806 | 0.1270 | -0.0647 | 0.0937 | -0.1011 |
| Aurora-B | 0.0853 | -0.1260 | -0.0297 | 0.0574 | 0.0044 | 0.0003 |
| MERIT40_pS29 | -0.0089 | 0.0987 | -0.0196 | 0.0035 | 0.0121 | -0.0276 |
| PRMT1 | 0.0701 | -0.0441 | -0.0488 | -0.0452 | 0.0399 | 0.1023 |
| Tau | 0.3797 | 0.2661 | -0.2984 | 0.0011 | -0.1063 | 0.0142 |
| NAPSIN-A | -0.0205 | 0.0651 | -0.0023 | 0.0097 | 0.0097 | -0.0408 |
| MITF | -0.0520 | -0.0184 | 0.1142 | 0.0768 | 0.0142 | -0.3204 |
| Akt2_pS474 | 0.0471 | 0.1593 | -0.2325 | -0.5014 | -0.0631 | 0.0998 |
| Akt1_pS473 | 0.0040 | 0.0223 | -0.0160 | -0.0705 | -0.0200 | 0.0351 |
| DM-Histone-H3 | -0.0121 | 0.0537 | -0.0035 | 0.0067 | -0.0288 | 0.0114 |
| Cox-IV | -0.0908 | 0.0516 | 0.1210 | -0.0542 | 0.0135 | -0.0088 |
| Glucocorticoid-Receptor | 0.0093 | -0.0216 | 0.5035 | -0.2032 | 0.3216 | -0.0917 |
| FAK | -0.0109 | -0.0988 | 0.0009 | 0.0065 | 0.0019 | -0.0462 |
| CD38 | -0.0892 | 0.1104 | -0.0620 | 0.0174 | -0.0146 | 0.0971 |
| AR | -0.0603 | -0.0068 | 0.0108 | 0.0634 | 0.0027 | -0.0466 |
| ADAR1 | 0.0432 | -0.0403 | -0.1260 | -0.1173 | 0.0361 | 0.1303 |
| Transglutaminase | 0.3232 | -0.3027 | -0.2285 | -0.1751 | 0.1779 | 0.2580 |
| Lasu1 | 0.0675 | -0.0769 | -0.0443 | 0.0402 | -0.0345 | 0.0392 |
| DNA_POLG | -0.0151 | -0.1007 | 0.0280 | 0.1012 | -0.0009 | -0.0668 |
| RSK1 | 0.0078 | -0.0085 | 0.0089 | -0.0388 | -0.0773 | 0.0506 |
| PMS2 | -0.0329 | -0.0021 | 0.0259 | 0.0086 | -0.0186 | 0.0515 |
| MACC1 | -0.0206 | -0.2303 | 0.2555 | 0.3410 | -0.0142 | 0.0171 |
| Pyk2_pY402 | -0.1762 | 0.0365 | -0.0360 | -0.1324 | 0.0471 | 0.0783 |
| p38-MAPK-_pT180_Y182 | -0.0585 | 0.0462 | -0.1497 | 0.1470 | 0.0642 | -0.1685 |
| S6_pS240_S244 | 0.1624 | -0.4324 | 0.3246 | -0.4706 | -0.0366 | 0.0413 |
| p27_pT198 | 0.0363 | 0.0099 | -0.1575 | -0.0137 | -0.0140 | 0.0326 |
| Collagen-VI | 0.0635 | -0.0188 | -0.1807 | -0.0819 | 0.0146 | 0.0937 |
| CRABP1 | -0.0458 | 0.1351 | -0.1157 | 0.0330 | -0.0302 | 0.1345 |
| PAICS | -0.4337 | -0.2521 | 0.6530 | 0.7201 | 0.0731 | -0.0684 |
| mTOR_pS2448 | -0.0155 | -0.0345 | 0.1578 | 0.0102 | 0.0826 | -0.1554 |
| INPP4b | -0.0598 | -0.0193 | 0.3957 | 0.0572 | 0.0151 | -0.0175 |
| Merlin | -0.0204 | 0.0080 | 0.1626 | -0.0619 | -0.0354 | 0.1093 |
| Gli3 | -0.1613 | 0.0789 | -0.0752 | 0.0825 | -0.1800 | 0.1604 |
| CD134 | -0.0164 | 0.0236 | -0.0083 | 0.0106 | -0.0078 | 0.0226 |
| HLA-DR-DP-DQ-DX | 0.0195 | -0.0356 | 0.0182 | -0.0057 | 0.0085 | -0.0418 |
| B-Raf_pS445 | -0.0163 | -0.0133 | 0.0049 | -0.0576 | 0.0817 | 0.0223 |
| N-Ras | -0.0245 | 0.0191 | -0.0338 | 0.0191 | -0.0381 | 0.0246 |
| p70-S6K1 | 0.0147 | -0.0347 | 0.0362 | -0.0464 | 0.0285 | -0.0181 |

| | | | | | | |
|---|---|---|---|---|---|---|
| ENY2 | 0.0941 | 0.0319 | -0.0595 | 0.0173 | -0.0292 | -0.0019 |
| XPF | -0.0047 | 0.0977 | -0.0067 | -0.0184 | -0.0388 | 0.0633 |
| Rheb | 0.6576 | -0.8928 | 0.8341 | -0.0609 | 0.0637 | -0.3695 |
| Connexin-43 | -0.2985 | 0.3659 | -0.0917 | 0.3807 | -0.0890 | 0.0937 |
| PKCa | -0.3207 | 0.3313 | 0.2202 | 0.2445 | -0.2235 | -0.2195 |
| Gys | -0.0044 | -0.0778 | 0.0127 | 0.0198 | -0.0115 | -0.0260 |
| AceCS1 | 0.0723 | -0.0312 | -0.0362 | -0.0119 | 0.0578 | 0.0272 |
| PKM2 | -0.0339 | 0.0216 | 0.1139 | -0.0679 | 0.1498 | -0.1449 |
| p38-a | 0.0893 | 0.0326 | -0.0322 | 0.0900 | -0.0395 | -0.0586 |
| ACSL1 | 0.1164 | -0.0337 | 0.0121 | -0.1344 | -0.0154 | 0.1066 |
| Aurora-A | 0.0987 | -0.0335 | -0.0034 | 0.0073 | -0.0045 | 0.1777 |
| VASP | 0.0146 | 0.0506 | -0.0635 | -0.0199 | 0.1892 | -0.0354 |
| CDK1_pT14 | -0.5758 | -0.0314 | 0.0469 | 0.0379 | -0.3554 | 0.1707 |
| ACLY_pS455 | 0.0026 | -0.0090 | -0.0849 | -0.0560 | 0.0048 | 0.0179 |
| FOXO3 | -0.0632 | 0.0515 | 0.0448 | 0.0533 | -0.0481 | -0.0549 |
| SOX7 | 0.0353 | 0.0090 | -0.0228 | -0.0115 | 0.0048 | -0.0001 |
| XIAP | 0.0066 | -0.0392 | 0.0714 | 0.0186 | -0.0158 | -0.0188 |
| HMHA1 | -0.0244 | 0.0948 | -0.0008 | -0.1014 | -0.0024 | 0.4477 |
| AMPKa | 0.1287 | -0.0300 | 0.0236 | -0.0163 | 0.0716 | -0.0205 |
| VEGFR2_pY1175 | -0.1060 | 0.0604 | -0.0057 | 0.0210 | 0.0016 | 0.0031 |
| FRS2-alpha_pY196 | -0.0161 | 0.0772 | -0.1212 | 0.1413 | -0.0786 | 0.0126 |
| Stathmin-1 | 0.0234 | -0.0277 | -0.0585 | 0.0342 | 0.0261 | -0.0622 |
| Cox2 | 0.0290 | 0.0199 | -0.0849 | -0.1300 | 0.0132 | -0.0085 |
| Akt1 | 0.0114 | -0.0426 | 0.0585 | -0.0278 | -0.0169 | 0.0217 |
| RRM2 | 0.0612 | -0.0350 | 0.0017 | 0.0057 | -0.0429 | 0.1756 |
| SHP2 | -0.0754 | 0.0777 | -0.0070 | 0.0707 | -0.0139 | 0.0163 |
| PAK1 | 0.0625 | 0.0071 | -0.0066 | -0.0275 | 0.0655 | -0.0337 |
| Cyclin-B1 | 0.0238 | -0.0289 | -0.0544 | 0.5907 | 0.0248 | -0.2843 |
| GATA3 | 0.1356 | -0.0045 | -0.0456 | -0.0505 | 0.0004 | 0.0434 |
| TTF1 | -0.0035 | 0.0168 | -0.0080 | -0.0707 | -0.0139 | 0.0405 |
| p90RSK_pT573 | -0.2574 | 0.2364 | 0.0520 | 0.1086 | -0.3196 | -0.0428 |
| DAPK1_pS308 | 0.1115 | -0.2395 | 0.1932 | 0.0186 | -0.0157 | -0.1482 |
| ATP5A | 0.0274 | -0.4899 | -0.0226 | 0.0300 | -0.2923 | 0.0691 |
| b-Catenin | 0.3741 | 0.0537 | -0.6353 | -0.9413 | 0.1377 | -0.0454 |
| ACC1 | -0.2855 | -0.0782 | 0.2478 | 0.1542 | 0.0741 | -0.0983 |
| KAP1 | 0.0315 | -0.0044 | -0.0415 | -0.0806 | 0.0003 | 0.0143 |
| TUFM | -0.0673 | -0.1079 | 0.1917 | 0.3024 | 0.0514 | -0.1183 |
| PLC-gamma2_pY759 | -0.1348 | 0.1308 | 0.0924 | -0.0037 | -0.0476 | 0.0190 |
| IMP3 | -0.0584 | 0.0401 | -0.0338 | 0.0631 | -0.0821 | 0.0430 |
| Rictor_pT1135 | -0.0513 | 0.0394 | 0.0051 | -0.0586 | -0.0053 | 0.0100 |
| Bak | 0.1545 | -0.0427 | -0.3606 | 0.0492 | 0.1465 | -0.1937 |
| mTOR | 0.0211 | -0.0836 | 0.1020 | -0.0265 | 0.0892 | -0.1386 |
| JNK_pT183_Y185 | -0.0352 | 0.1654 | 0.0238 | 0.1864 | -0.0419 | -0.1461 |
| ULK1_pS757 | -0.0336 | 0.0568 | 0.0222 | -0.0437 | -0.0514 | 0.0747 |
| eIF4E | -0.1235 | 0.0171 | -0.0167 | -0.1123 | 0.0281 | 0.0504 |
| Chk2 | 0.1977 | -0.1148 | -0.1423 | -0.0800 | 0.0828 | 0.1009 |
| UBQLN4 | -0.0039 | 0.0128 | -0.0392 | 0.0000 | 0.0028 | -0.0120 |
| MR1 | 0.0164 | 0.0304 | -0.0365 | -0.0256 | -0.0115 | 0.0162 |
| PYGM | 0.0244 | -0.0003 | -0.0398 | 0.0038 | -0.0035 | 0.0087 |
| Rad51 | -0.0061 | 0.0033 | -0.0028 | 0.0183 | -0.0272 | -0.0147 |
| D-a-Tubulin | -0.0299 | 0.0503 | 0.0184 | -0.0237 | -0.0465 | 0.0583 |

| | | | | | | |
|---|---|---|---|---|---|---|
| EGFR | -0.1145 | -0.0090 | 0.3205 | 0.0722 | -0.0692 | 0.0173 |
| Chk1 | -0.0013 | 0.0344 | -0.0471 | -0.0129 | -0.0147 | 0.0562 |
| CD45 | 0.0615 | -0.0409 | 0.0262 | 0.0029 | -0.0075 | 0.0122 |
| S6 | 0.2559 | 0.0933 | -0.0955 | -0.0869 | 0.2789 | -0.1519 |
| YB1_pS102 | 0.0000 | -0.0588 | 0.0547 | 0.0947 | -0.0316 | -0.0034 |
| Gab2 | -0.0330 | 0.0933 | -0.0232 | 0.0315 | -0.0988 | 0.0295 |
| PDHK1 | -0.0364 | 0.0306 | 0.0216 | -0.0178 | -0.0094 | 0.0141 |
| Glutaminase | -0.0162 | -0.0490 | 0.0796 | -0.0093 | 0.0002 | 0.0847 |
| eEF2K | 0.0214 | -0.0068 | -0.0285 | -0.1091 | 0.0027 | 0.0350 |
| Bax | 0.1266 | 0.0423 | -0.1902 | 0.0031 | -0.0003 | -0.1566 |
| UGT1A | 0.2957 | 0.0141 | -0.2761 | -0.2178 | 0.0349 | -0.0057 |
| LDHA | 0.0977 | -0.3283 | 0.0305 | -0.0231 | 0.1237 | -0.1214 |
| Dvl3 | 0.1306 | -0.1262 | -0.0479 | -0.1176 | 0.1451 | 0.0571 |
| CDT1 | -0.0119 | 0.1994 | -0.0824 | 0.0066 | -0.0425 | 0.0378 |
| Myosin-IIa_pS1943 | 0.0934 | -0.1011 | -0.0162 | -0.0529 | 0.0841 | 0.0254 |
| CD74 | 0.0748 | 0.0696 | -0.1281 | -0.0713 | -0.0225 | 0.0272 |
| Bim | -0.0049 | 0.0025 | -0.1583 | -0.1438 | 0.0711 | 0.0058 |
| Smac | 0.1084 | -0.1512 | 0.0448 | 0.1343 | -0.2509 | -0.0355 |
| DNA-Ligase-IV | -0.1058 | -0.0470 | 0.1654 | 0.2142 | 0.0428 | -0.1061 |
| MTCO1 | 0.0807 | -0.1959 | 0.2062 | -0.0861 | 0.1109 | -0.0846 |
| p27_pT157 | 0.0122 | 0.0010 | -0.0006 | 0.0322 | -0.0298 | -0.0104 |
| SLC1A5 | 0.0061 | -0.2501 | 0.1750 | 0.1386 | -0.0221 | -0.0377 |
| Cadherin-6 | 0.0085 | -0.0208 | 0.1178 | 0.1508 | -0.0901 | -0.0476 |
| cdc25C | 0.0760 | -0.0170 | -0.0831 | -0.0554 | 0.0129 | 0.0844 |
| Enolase-1 | -0.0780 | -0.0421 | 0.0975 | 0.1194 | 0.0380 | -0.3432 |
| MERIT40 | 0.0208 | -0.0516 | 0.0012 | -0.0953 | 0.0132 | 0.0081 |
| PREX1 | -0.1680 | 0.1334 | -0.0136 | 0.0218 | 0.0103 | -0.2008 |
| Bad_pS112 | 0.0057 | -0.0642 | 0.0217 | 0.0231 | -0.0202 | -0.0819 |
| ER-a | 0.0202 | 0.2101 | -0.0667 | 0.1278 | -0.0362 | -0.0300 |
| PI3K-p110-b | -0.0090 | 0.0220 | -0.0025 | 0.0570 | -0.0305 | -0.0516 |
| FoxO3a_pS318_S321 | -0.0036 | 0.0207 | -0.0094 | -0.0126 | 0.0386 | 0.0001 |
| NDUFB4 | -0.0095 | 0.0075 | -0.0034 | 0.0041 | -0.0652 | 0.0271 |
| ASNS | -0.0742 | -0.2019 | 0.4201 | 0.4160 | -0.0963 | 0.0708 |
| cdc2_pY15 | 0.0761 | 0.0459 | -0.0454 | -0.1128 | -0.0907 | 0.1786 |
| AMPK-a2_pS345 | -0.0335 | 0.0123 | 0.0764 | -0.0058 | -0.0195 | 0.0435 |
| DYRK1B | 0.0377 | 0.0192 | -0.0505 | -0.0269 | 0.0104 | -0.0057 |
| Rad23A | 0.0077 | 0.0522 | -0.1627 | -0.1999 | -0.0173 | 0.0220 |
| BMK1-Erk5_pT218_Y220 | 0.0214 | -0.0349 | 0.0375 | 0.1086 | -0.0374 | -0.1137 |
| TRIM25 | -0.1582 | -0.0773 | 0.2198 | 0.0838 | 0.2557 | -0.1886 |
| PARP | 0.0640 | -0.0134 | -0.0117 | -0.0520 | 0.0085 | 0.1429 |
| 4E-BP1_pS65 | 0.0473 | -0.3217 | 0.3727 | 0.1825 | -0.0633 | -0.2441 |
| FGFR1 | -0.0911 | 0.0653 | -0.0113 | 0.0830 | -0.0801 | 0.0206 |
| Bcl-xL | -0.0240 | -0.1966 | 0.0337 | 0.0252 | 0.0080 | -0.1400 |
| VHL-EPPK1 | 0.1089 | 0.0256 | -0.2236 | -0.2047 | -0.0027 | 0.0074 |
| Paxillin | 0.0076 | -0.0304 | -0.0190 | -0.1116 | 0.0650 | 0.0578 |
| Beclin | -0.0327 | 0.0043 | 0.0123 | 0.0465 | -0.0408 | 0.0040 |
| Chk1_pS345 | 0.1062 | -0.0571 | -0.0120 | 0.0053 | 0.0692 | 0.0100 |
| ACC_pS79 | 0.2175 | -1.1031 | -0.0407 | 0.0481 | 0.1500 | -0.7025 |
| HSP27_pS82 | 0.0147 | -0.0812 | -0.0086 | 0.0544 | 0.0054 | -0.2084 |
| H2AX_pS140 | 0.0024 | -0.0718 | 0.0274 | 0.0342 | -0.0184 | -0.0278 |
| 53BP1 | -0.0469 | -1.9282 | 0.2355 | 0.0416 | 0.1668 | -1.2666 |

| | | | | | | |
|---|---|---|---|---|---|---|
| Creb | -0.0114 | -0.0259 | 0.0952 | 0.0430 | -0.0045 | -0.0155 |
| Midkine | -0.0912 | -0.0175 | 0.1399 | 0.1551 | 0.0134 | -0.0606 |
| Sox2 | -0.1926 | 0.0104 | 0.0528 | -0.0855 | -0.0145 | 0.0397 |
| IGFRb | 0.4952 | 0.0285 | -0.0281 | 0.5409 | -0.0742 | -0.2047 |
| 14-3-3-epsilon | -0.1006 | 0.0622 | 0.0488 | 0.1118 | -0.0521 | -0.1204 |
| Chk1_pS296 | -0.0703 | 0.0683 | 0.0081 | -0.0745 | -0.0113 | 0.1571 |
| PCNA | 0.0923 | -0.0698 | 0.0205 | -0.0934 | 0.0904 | -0.0112 |
| EGFR_pY1173 | 0.1964 | -0.3214 | 0.1352 | 0.0167 | -0.0138 | -0.1290 |
| Rb | 0.0209 | -0.1338 | 0.0445 | 0.0756 | -0.0368 | -0.1807 |
| MLH1 | 0.0668 | -0.1286 | 0.6566 | 0.5162 | -0.0828 | -0.0825 |
| Caspase-3-cleaved | -0.0217 | 0.0156 | 0.0103 | 0.0659 | -0.0311 | -0.0064 |
| PEA-15_pS116 | 0.0033 | 0.0797 | -0.0569 | 0.0714 | -0.0193 | -0.0694 |
| Atg7 | 0.0423 | 0.0087 | -0.0083 | -0.0111 | 0.0953 | -0.0127 |
| YES1 | -0.0137 | -0.0008 | 0.0013 | 0.0341 | 0.0077 | -0.0389 |
| Pdcd4 | -0.2600 | 0.5030 | 0.1742 | 0.7490 | -0.4817 | -0.1649 |
| XBP-1 | 0.0973 | -0.2038 | 0.6628 | 0.5409 | -0.1290 | -0.1007 |
| MIF | -0.6339 | 0.0741 | 0.1473 | -0.0398 | 0.0426 | -0.2267 |
| Aurora-ABC_pT288_pT232_pT198 | -0.1891 | 0.0065 | 0.0203 | 0.0000 | -0.1282 | 0.0246 |
| VEGFR-2 | 0.0438 | -0.0166 | -0.2029 | -0.1880 | 0.0124 | 0.0755 |
| Snail | 0.4314 | -0.0297 | -0.2273 | -0.2991 | 0.2592 | 0.0380 |
| Histone-H3 | 0.1142 | -0.0182 | -0.1333 | -0.1564 | 0.0140 | 0.0425 |
| UQCRC2 | -0.1372 | 0.1031 | 0.0282 | 0.0997 | -0.0324 | -0.0189 |
| PHLPP | -0.0133 | -0.0097 | 0.0449 | 0.0079 | -0.0569 | 0.0477 |
| b-Catenin_pT41_S45 | -0.0123 | -0.0001 | -0.0064 | 0.0113 | 0.0008 | -0.0238 |
| FGF-basic | -0.0657 | -0.0660 | 0.1004 | 0.0770 | 0.0497 | -0.1065 |
| Granzyme-B | -0.0603 | 0.1275 | 0.0489 | -0.2208 | -0.1088 | 0.6164 |
| PIP4K2A | -0.1407 | 0.0666 | 0.1130 | 0.2026 | -0.1934 | -0.0582 |
| BCL2A1 | 0.0539 | 0.0774 | -0.1264 | 0.2838 | -0.1319 | -0.0574 |
| HER3_pY1289 | 0.0199 | -0.1555 | 0.0519 | 0.0017 | 0.0011 | -0.1359 |
| PDHA1 | 0.0349 | -0.0750 | -0.0053 | 0.0160 | 0.0021 | -0.0410 |
| PI3K-p110-a | 0.0167 | 0.1567 | -0.0450 | 0.2008 | -0.1202 | -0.0202 |
| CD2 | 0.0195 | 0.0544 | -0.0506 | 0.0001 | 0.0023 | 0.0024 |
| 4E-BP1 | 0.0783 | -0.0939 | 0.0397 | 0.0095 | -0.0224 | 0.0058 |
| ATP5H | 0.0443 | -0.0019 | 0.0023 | -0.0244 | -0.0171 | 0.1140 |
| Jagged1 | -0.0381 | 0.0231 | 0.0449 | -0.0166 | -0.0292 | 0.0552 |
| VAV1 | -0.0302 | -0.0291 | 0.8315 | 1.0127 | 0.0126 | -0.0079 |
| Hif-1-alpha | -0.0232 | 0.0034 | 0.0593 | 0.0798 | -0.0076 | -0.0117 |
| HSP60 | -0.1210 | 0.0024 | 0.1211 | 0.1029 | -0.0066 | -0.0484 |
| PKC-b-II_pS660 | -0.1444 | 0.0356 | 0.0685 | -0.0389 | -0.0397 | 0.0802 |
| TFAM | -0.0987 | 0.1323 | -0.0194 | 0.0268 | 0.1440 | -0.1032 |
| c-Myc | 0.1889 | 0.0478 | -0.3518 | -0.0273 | -0.0439 | 0.0426 |
| Caspase-7-cleaved | 0.0083 | 0.0870 | -0.1242 | 0.1256 | -0.0243 | -0.0847 |
| Src_pY416 | -0.0403 | 0.2147 | -0.1978 | -0.1074 | 0.1171 | 0.0368 |
| MTSS1 | -0.0150 | 0.0454 | -0.0287 | 0.0097 | -0.0343 | 0.0124 |
| YAP_pS127 | 0.0849 | 0.0260 | -0.1113 | -0.0901 | 0.1545 | -0.0177 |
| PRC1_pT481 | -0.0210 | 0.0592 | 0.0096 | 0.1834 | -0.0912 | -0.0018 |
| Rad50 | 0.0241 | -0.0384 | 0.0468 | -0.0294 | 0.0705 | -0.0296 |
| WIPI1 | 0.1345 | -0.0321 | -0.1153 | -0.1342 | 0.0345 | 0.0405 |
| Ets-1 | -0.0267 | 0.1289 | -0.2085 | -0.2077 | 0.0107 | 0.0298 |
| IRF-3 | 0.0295 | 0.0035 | -0.0357 | 0.0030 | 0.0577 | -0.0859 |
| TRAP1 | -0.0505 | -0.0300 | 0.5401 | 0.6414 | -0.0789 | 0.0383 |

| | | | | | | |
|---|---|---|---|---|---|---|
| Caveolin-1 | 0.0410 | 0.1604 | -0.1557 | -0.0464 | 0.1574 | -0.1236 |
| CHD1L | 0.0516 | -0.0105 | -0.0235 | -0.0120 | 0.0063 | 0.0359 |
| Hexokinase-I | -0.0442 | 0.0615 | 0.0145 | 0.1208 | -0.0567 | -0.0052 |
| Slfn11 | 0.0283 | -0.2808 | 0.2584 | 0.1341 | -0.0443 | -0.1612 |
| Elk1_pS383 | 0.0517 | 0.0175 | -0.1537 | 0.0567 | -0.0217 | -0.0874 |
| PLC-gamma1 | -0.0013 | -0.1645 | 0.5465 | 0.5465 | -0.0428 | -0.0022 |
| TROP2 | -0.0375 | -0.0838 | 0.3104 | -0.1225 | 0.2131 | 0.0340 |
| HLA-DQA1 | -0.0011 | 0.0064 | -0.0510 | -0.0632 | -0.0106 | 0.0162 |
| RRM1 | -0.0122 | 0.0750 | -0.1119 | -0.0725 | -0.0038 | 0.0525 |
| Puma | 0.0195 | 0.0036 | -0.1194 | -0.1723 | -0.0077 | 0.1204 |
| MEK2 | 0.0347 | -0.0021 | -0.0203 | 0.0086 | 0.0122 | -0.0022 |
| Mitofusin-2 | 0.1940 | -0.0151 | 0.0155 | -0.0418 | 0.2143 | -0.0169 |
| CD171 | -0.1172 | -0.1348 | 0.3522 | 0.3738 | 0.1012 | -0.4848 |
| SF2 | 0.0025 | 0.0448 | -0.0816 | -0.0309 | -0.0185 | 0.0294 |
| B7-H4 | -0.0693 | 0.0295 | 0.0109 | -0.0035 | -0.0801 | 0.0394 |
| Akt_pT308 | -0.0433 | 0.0276 | -0.0014 | -0.0650 | 0.0502 | 0.0107 |
| PRAS40_pT246 | 0.0471 | -0.2390 | 0.2149 | 0.3692 | -0.0630 | -0.3637 |
| Notch3 | -0.0085 | 0.0322 | -0.0133 | 0.0056 | -0.0028 | 0.0030 |
| Annexin-I | -0.0053 | -0.0314 | 0.1188 | 0.0951 | -0.0107 | -0.0194 |
| ZAP-70 | -0.0058 | 0.0650 | -0.0142 | 0.0005 | -0.0234 | 0.0580 |
| Caspase-8 | 0.1592 | -0.0408 | -0.1123 | -0.1902 | 0.1132 | 0.0491 |
| GATA6 | -0.1901 | 0.1094 | 0.0599 | -0.0525 | -0.1227 | 0.1901 |
| N-Cadherin | -0.0263 | -0.0308 | 0.0886 | 0.0605 | -0.0662 | 0.0228 |
| PAK_pT423_T402 | -0.0325 | -0.0279 | 0.0306 | 0.0271 | 0.0238 | -0.0900 |
| Tuberin_pT1462 | -0.0772 | 0.1073 | -0.0431 | 0.0642 | 0.0399 | -0.1346 |
| YTHDF2 | -0.0109 | -0.0513 | 0.3854 | 0.3455 | -0.0051 | -0.0538 |
| Src | -0.1214 | -0.1038 | 0.3488 | 0.2392 | 0.0996 | -0.1781 |
| p53 | -0.0932 | 0.0261 | 0.0119 | -0.0524 | -0.0116 | 0.0163 |
| B7-H3 | -0.4056 | 0.1524 | 0.1135 | -0.1136 | -0.0059 | 0.0106 |
| CDK9 | -0.0040 | 0.0168 | -0.0350 | 0.0024 | 0.0004 | -0.0279 |
| IR-b | 0.0077 | -0.0739 | -0.0028 | 0.0102 | 0.0594 | -0.1293 |
| HER2 | -0.0190 | -0.1711 | 0.5241 | 0.4493 | -0.0927 | 0.0156 |
| Rab11 | -0.0526 | 0.0457 | 0.0145 | -0.0011 | -0.0094 | 0.0141 |
| PDH | -0.0156 | 0.0041 | 0.0034 | 0.0072 | -0.0044 | -0.0099 |
| Porin | 0.0318 | 0.0171 | -0.0725 | -0.0374 | -0.0213 | 0.0302 |
| METTL3 | 0.0023 | 0.0436 | -0.0720 | -0.0439 | -0.0183 | 0.0309 |
| PD-L1 | -0.1975 | 0.2806 | 0.0437 | -0.0363 | -0.1173 | 0.2596 |
| PRAS40 | 0.0159 | 0.0486 | -0.0597 | 0.0017 | -0.0142 | 0.0136 |
| Patched | -0.0188 | 0.0184 | 0.0074 | -0.0200 | -0.0136 | 0.0430 |
| HES1 | -0.1376 | 0.0253 | 0.0518 | -0.0588 | -0.0050 | 0.0097 |
| FN14 | 0.1346 | -0.0530 | -0.0743 | -0.1517 | 0.0488 | 0.1004 |
| JNK2 | 0.1388 | 0.2871 | -0.8216 | 0.4426 | -0.7428 | -0.1423 |
| CD5 | 0.0333 | 0.0343 | -0.0603 | 0.0000 | -0.0089 | 0.0136 |
| Ambra1_pS52 | 0.0599 | -0.0371 | -0.0161 | 0.0298 | 0.0129 | -0.0381 |
| XRCC1 | -0.0665 | 0.2188 | -0.0669 | 0.0856 | -0.0509 | 0.0556 |
| c-IAP2 | 0.0322 | -0.0790 | 0.0329 | -0.0614 | 0.0700 | -0.0236 |
| CDKN2A | -1.0431 | 0.2940 | 0.4753 | 0.2501 | -0.2808 | -0.2347 |
| Shc_pY317 | 0.1543 | 0.3053 | -0.8832 | 0.5632 | -0.8241 | -0.1578 |
| S6_pS235_S236 | 0.0111 | -0.1261 | 0.1900 | -0.0164 | 0.1414 | -0.0503 |
| U-Histone-H2B | 0.0437 | -0.0070 | -0.0237 | -0.0356 | 0.0029 | 0.0458 |

| | | | | | | |
|---|---|---|---|---|---|---|
| | **OV7** | | | | | |
| | **DMF/MCT** | **DMF/MCT** | **DMF/MCT** | **6 uM Broad Spectrum CBD** | **6 uM Broad Spectrum CBD** | **6 uM Broad Spectrum CBD** |
| Wee1 | 0.0727 | 0.0900 | -0.0565 | 0.0238 | 0.0171 | -0.0102 |
| ATM | 0.2639 | 0.2146 | -0.0423 | 0.0630 | -0.0658 | -0.0448 |
| LCN2 | 0.0056 | -0.0397 | 0.0307 | 0.0130 | -0.0050 | 0.1001 |
| Lyn | 0.1382 | 0.0008 | 0.0763 | 0.0237 | -0.1139 | -0.1392 |
| CENP-A | 0.0404 | 0.0213 | 0.0425 | -0.3586 | -0.0153 | -0.1229 |
| ERCC5 | 0.0718 | 0.0276 | -0.1060 | -0.1220 | -0.0216 | 0.0998 |
| Glutamate-D1-2 | 0.0991 | -0.0526 | -0.1270 | 0.1531 | -0.2026 | 0.0508 |
| Grp75 | -0.0008 | -0.0150 | 0.2499 | 0.2613 | -0.1351 | -0.2250 |
| RIP | 0.0414 | 0.0975 | -0.0261 | 0.0468 | -0.0699 | -0.1311 |
| IRF-1 | 0.0330 | 0.0372 | -0.0166 | -0.1776 | 0.0188 | -0.2501 |
| G6PD | 0.0142 | 0.0224 | 0.0867 | -0.0489 | -0.0366 | -0.0214 |
| E2F1 | 0.0239 | 0.0593 | -0.0435 | 0.1330 | -0.0745 | -0.0711 |
| SOD2 | 0.1271 | -0.0363 | 0.0083 | 0.2304 | -0.1573 | -0.0139 |
| ERCC1 | 0.0295 | -0.0727 | 0.0517 | -0.0875 | -0.0290 | 0.1007 |
| Folliculin | -0.0203 | -0.0052 | 0.0181 | -0.3604 | 0.1393 | 0.0034 |
| eIF4G | 0.0533 | 0.1253 | 0.1534 | -0.1212 | -0.0528 | -0.0794 |
| S100A4 | 0.0239 | 0.0174 | 0.0650 | -0.1615 | -0.0161 | -0.0192 |
| ALKBH5 | 0.2276 | 0.1472 | 0.0398 | -0.5305 | -0.0377 | -0.5804 |
| c-Abl_pY412 | -0.0051 | -0.0107 | 0.0531 | -0.5676 | -0.0193 | 0.0215 |
| Cyclin-D1 | 0.0086 | -0.0146 | 0.0004 | -0.1264 | -0.0057 | 0.0593 |
| RPA32_pS4_S8 | -0.0175 | 0.0230 | -0.0427 | 0.0754 | 0.0181 | -0.0644 |
| CREB_pS133 | 0.0873 | 0.1396 | 0.1937 | -0.3295 | -0.0868 | -0.1178 |
| PKA-a | 0.7856 | -0.0341 | -0.1933 | 0.2045 | -0.1549 | 0.0323 |
| Synaptophysin | 0.0811 | 0.0256 | 0.1686 | -0.0011 | -0.1212 | -0.0466 |
| HSP70 | 0.0039 | -0.0197 | -0.0456 | 0.1077 | 0.0309 | -0.0302 |
| p21 | 0.0083 | 0.0150 | 0.1065 | -0.0317 | -0.0096 | -0.0155 |
| YTHDF3 | 0.0155 | 0.0168 | 0.0181 | -0.2856 | -0.0109 | -0.1448 |
| ATM_pS1981 | -0.0121 | -0.0773 | 0.5706 | 0.2975 | -0.0329 | 0.0049 |
| RBM15 | -0.0506 | 0.0348 | 0.0996 | -0.2197 | -0.0307 | 0.1068 |
| MEK1 | -0.4502 | 0.3825 | 0.1572 | 0.0126 | 0.0283 | -0.1549 |
| FANCD2 | 0.0192 | -0.0350 | 0.0735 | -0.0259 | -0.0439 | 0.0827 |
| FASN | 0.0968 | 0.0128 | 0.4041 | -0.9066 | -0.0069 | -0.2569 |
| Rb_pS807_S811 | 0.0901 | 0.1011 | -0.0997 | -1.1232 | 0.1018 | -0.6645 |
| PAR | -0.1347 | 0.9598 | 0.9241 | -0.1250 | -0.3035 | 0.1275 |
| Notch1 | 0.0001 | -0.0185 | 0.0741 | -0.0989 | 0.0005 | 0.0163 |
| SOD1 | -0.0454 | -0.0786 | -0.0790 | 0.2154 | 0.0460 | 0.0579 |
| Stat5a | 0.0790 | 0.0917 | -0.0111 | -1.9464 | 0.0132 | -0.3104 |
| Myt1 | 0.0012 | 0.1241 | -0.0828 | 0.1550 | -0.0007 | -0.1593 |
| eIF4E_pS209 | 0.0181 | 0.0641 | -0.0161 | -0.0211 | 0.0182 | -0.0753 |
| PKC-a-b-II_pT638_T641 | 0.0529 | 0.2332 | -0.4228 | -0.3243 | 0.0797 | -0.0601 |
| ATR_pS428 | -0.1381 | 0.1541 | 0.0176 | 0.0031 | 0.0569 | -0.0994 |
| Sox17 | -0.0449 | -0.0086 | 0.1174 | 0.1240 | -0.0328 | 0.0068 |
| TSC1 | 0.0341 | -0.0088 | -0.0374 | 0.0701 | -0.0435 | 0.0070 |
| BRD4 | 0.1487 | 0.0328 | -0.0470 | -0.6037 | 0.0905 | -0.6027 |
| DJ1 | 0.1643 | 0.0235 | -0.0377 | 0.1171 | -0.1092 | -0.2105 |
| Akt2 | 0.0071 | 0.0666 | -0.0006 | -0.0735 | 0.0027 | -0.1336 |

| | | | | | | |
|---|---|---|---|---|---|---|
| CD44 | 0.2091 | 0.0616 | 0.1838 | -0.3081 | -0.1517 | -0.0634 |
| YAP | -0.0762 | -0.3359 | 0.0565 | -0.5354 | 0.1084 | 0.2051 |
| c-Jun_pS73 | -0.0091 | 0.0509 | 0.3519 | -0.0227 | -0.0853 | 0.0018 |
| Erk5 | 0.0026 | -0.0184 | 0.5507 | -1.6481 | -0.0139 | 0.2645 |
| IRS1 | 0.0211 | -0.0529 | 0.3728 | 0.5809 | -0.5262 | -0.0283 |
| EphA2_pS897 | -0.2179 | 0.4186 | 0.3391 | 0.0505 | -0.0712 | -0.0174 |
| CIITA | 0.0099 | -0.0008 | -0.0139 | 0.0263 | 0.0006 | 0.0068 |
| PHGDH | -0.0052 | -0.0101 | -0.1968 | 0.0549 | 0.0161 | -0.1434 |
| Src_pY527 | -0.0292 | 0.1566 | 0.1384 | -0.8688 | 0.0298 | -0.3643 |
| ZEB1 | 0.1478 | 0.2140 | 0.2304 | -0.1506 | -0.1569 | -0.1550 |
| Cyclin-D3 | -0.2082 | 0.4901 | 0.5495 | 0.2273 | -0.2122 | -0.3260 |
| MSI2 | -0.0017 | 0.0370 | -0.0179 | -0.1869 | 0.0226 | -0.0272 |
| Cdc6 | 0.0223 | -0.0381 | 0.3395 | -0.4996 | -0.1070 | 0.0494 |
| IDO | -0.0095 | -0.0739 | 0.2198 | 0.1645 | -0.0439 | 0.0022 |
| EphA2_pY588 | 0.0534 | -0.0348 | 0.1392 | -2.5002 | 0.0303 | -0.0157 |
| NF-kB-p65_pS536 | 0.2448 | 0.2655 | -0.1886 | -0.1572 | 0.0877 | -0.0732 |
| ATR | -0.0752 | -0.1354 | -0.0172 | 0.2066 | 0.0193 | 0.0403 |
| P-Cadherin | -0.0016 | 0.0282 | -0.0321 | -0.0380 | 0.0195 | -0.0050 |
| p70-S6K_pT389 | -0.0068 | 0.0601 | -0.2738 | -0.3634 | 0.0074 | 0.0177 |
| Wee1_pS642 | 0.0961 | 0.1033 | 0.0010 | -0.2289 | 0.0011 | -0.0883 |
| Akt | 0.1012 | 0.1138 | 0.0731 | -0.2514 | -0.0710 | -0.3747 |
| Hexokinase-II | 0.2474 | 0.0781 | -0.0747 | -0.2944 | 0.0768 | -0.5189 |
| Melanoma-gp100 | -0.0469 | -0.0915 | 0.0288 | 0.1495 | -0.0363 | 0.0397 |
| CtIP | 0.0204 | -0.0047 | 0.0847 | -0.4169 | -0.1240 | 0.0029 |
| IGFBP2 | -0.2237 | 0.0607 | 0.0006 | 0.0201 | 0.1300 | -0.1929 |
| Stat1_pY701 | -0.0040 | -0.0296 | -0.0057 | 0.0267 | -0.0549 | 0.0064 |
| CD20 | 0.0296 | -0.0454 | 0.1348 | -0.2283 | -0.1222 | 0.0926 |
| TRIM24 | -0.0009 | -0.0776 | 0.0292 | -0.0570 | 0.0014 | 0.1029 |
| CA9 | -0.0448 | -0.0127 | -0.0343 | 0.0372 | 0.0416 | 0.0151 |
| ERRalpha | -0.0384 | 0.0226 | 0.1283 | -1.5945 | 0.0742 | -0.4518 |
| PGM1 | 0.0990 | 0.0473 | -0.0448 | -0.4262 | 0.0470 | -0.0541 |
| CD31 | -0.0509 | -0.0004 | 0.1068 | -0.7151 | 0.1504 | -0.0014 |
| UBAC1 | -0.0582 | 0.0630 | 0.0385 | -0.0827 | -0.0510 | 0.0676 |
| BRCA1 | -0.0054 | -0.0247 | 0.0109 | 0.0245 | 0.0054 | 0.1294 |
| DDR1_pY513 | -0.0056 | -0.0102 | 0.0210 | -0.1507 | -0.0682 | 0.0385 |
| PIP4K2B | -0.0168 | -0.0012 | 0.0291 | 0.0914 | 0.0072 | -0.0522 |
| Folate-Binding-Protein | -0.0904 | -0.0122 | 0.0740 | 0.0793 | 0.0182 | -0.0037 |
| TFRC | -0.0337 | 0.1878 | 0.0141 | -0.2393 | -0.0217 | 0.0289 |
| PDK1_pS241 | 0.2706 | 0.2246 | 0.0888 | -0.0681 | -0.1833 | -0.4598 |
| JAB1 | -0.0281 | -0.0835 | -0.0286 | 0.2844 | 0.0286 | 0.1787 |
| Annexin-VII | 0.0099 | 0.0210 | -0.1171 | 0.1195 | -0.0094 | -0.1104 |
| c-Kit | 0.4572 | 0.4617 | -0.6877 | 0.0202 | 0.0206 | -0.3296 |
| CD4 | -0.0087 | 0.0556 | 0.1287 | -0.4914 | 0.0081 | 0.0015 |
| DM-K9-Histone-H3 | -0.0064 | -0.0098 | 0.0258 | -0.1199 | 0.0114 | 0.0031 |
| Rab11FIP1 | 0.0333 | 0.3906 | -0.0530 | -0.3414 | 0.0689 | -0.5197 |
| MSH6 | 0.0635 | 0.2739 | 0.0191 | -0.3182 | -0.0926 | -0.0248 |
| PTPN12 | 0.0955 | 0.0741 | 0.1290 | -0.0496 | -0.1641 | -0.1884 |
| Myosin-IIa | -0.1909 | 0.4165 | 0.3502 | -0.1615 | 0.1440 | -0.1295 |
| LRP6_pS1490 | -0.0007 | 0.0078 | -0.0438 | 0.0205 | 0.0153 | -0.0007 |
| PDK1 | 0.1230 | 0.1780 | -0.0200 | -0.0473 | 0.0221 | -0.2766 |
| PAK_pS474_S602_S560 | 0.1290 | 0.1513 | 0.0358 | -0.2400 | -0.0480 | -0.0415 |

| | | | | | | |
|---|---|---|---|---|---|---|
| c-Met_pY1234_Y1235 | -0.0345 | -0.0657 | -0.0634 | 0.2991 | 0.0350 | 0.0656 |
| MYH11 | -0.0561 | 0.0467 | -0.1708 | 0.0588 | 0.0342 | -0.0197 |
| ARID1A | -0.0734 | 0.0557 | 0.0333 | -0.1555 | 0.0853 | -0.0389 |
| H2AX_pS139 | -0.0783 | 0.1315 | -0.0611 | 0.0818 | 0.0740 | -0.1421 |
| Rictor | 0.0265 | 0.0019 | -0.0161 | 0.0406 | -0.1313 | -0.0739 |
| UVRAG | 0.0086 | -0.0084 | -0.0601 | -0.2073 | 0.0439 | 0.0066 |
| p27-Kip1 | 0.1104 | -0.0493 | 0.0215 | -0.0764 | -0.0194 | 0.2215 |
| TEAD | -0.0888 | 0.0028 | 0.1594 | 0.0622 | -0.1413 | -0.0046 |
| A-Raf | 0.0352 | -0.1602 | -0.0549 | 0.7652 | 0.1610 | -0.7834 |
| Cdc42 | 0.0262 | 0.0044 | -0.0853 | -0.2506 | 0.0134 | 0.0011 |
| NDRG1_pT346 | 0.2081 | 0.1467 | 0.1501 | -0.2427 | -0.1407 | -0.4606 |
| B-Raf | 0.1062 | 0.2216 | -0.4446 | -0.2948 | 0.0905 | -0.0760 |
| SIRP-alpha | 0.2228 | 0.1039 | -0.5556 | -0.3843 | 0.2470 | -0.1057 |
| ATRX | -0.0591 | -0.0104 | -0.1195 | 0.1941 | 0.4066 | 0.0086 |
| A-Raf_pS299 | -0.0158 | -0.1461 | 0.1143 | -0.1442 | 0.0163 | 0.0577 |
| MEK1_pS217_S221 | 0.1263 | 0.2117 | 0.0966 | -0.0759 | -0.1703 | -0.2966 |
| E-Cadherin | -0.0194 | -0.0093 | 0.2708 | -0.2653 | 0.1441 | 0.0075 |
| SGK1 | 0.1242 | 0.0188 | 0.1200 | 0.0057 | -0.1290 | -0.0280 |
| SDHA | -0.0090 | -0.1053 | -0.1324 | 0.4397 | 0.0095 | 0.2794 |
| GCN5L2 | -0.0679 | -0.0253 | -0.0761 | 0.3376 | 0.0312 | 0.3777 |
| p44-42-MAPK | 0.1408 | 0.1456 | 0.0128 | -0.2137 | -0.0107 | -0.5759 |
| MIG6 | -0.1798 | -0.3829 | 0.1160 | 0.2984 | 0.1490 | -0.1216 |
| RSK | 0.0422 | -0.0580 | 0.0906 | -0.1489 | -0.0777 | 0.1846 |
| PLK1 | 0.1446 | 0.2692 | 0.5720 | -0.9936 | -0.1642 | -0.1518 |
| KEAP1 | 0.0337 | 0.0540 | 0.0900 | -0.2437 | -0.0332 | -0.2079 |
| MLKL | 0.0124 | 0.0745 | -0.0252 | -0.0167 | 0.0273 | -0.1393 |
| ACVRL1 | -0.0150 | 0.0181 | 0.0567 | -0.4693 | 0.0156 | -0.0478 |
| WIPI2 | -0.0245 | 0.0829 | 0.0049 | -0.4948 | 0.0885 | -0.0344 |
| Caspase-8-cleaved | -0.0097 | -0.0676 | -0.0100 | 0.1233 | 0.0021 | 0.0394 |
| Akt_pS473 | 0.2456 | 0.1219 | 0.1167 | -1.4661 | -0.1146 | -0.3875 |
| DUSP6 | -0.1443 | -0.3320 | 0.1247 | 0.7661 | -0.3963 | 0.1486 |
| HSP27 | -0.0953 | -0.1496 | 0.0485 | 0.1251 | -0.0464 | 0.0731 |
| HER2_pY1248 | -0.0001 | -0.0088 | 0.0920 | -0.4612 | -0.0282 | 0.0070 |
| LAD1 | 0.1551 | -0.0722 | 0.0253 | -0.3137 | 0.3567 | -0.0309 |
| c-Abl | 0.0015 | -0.0131 | -0.1915 | 0.0377 | 0.1897 | -0.1011 |
| TAZ | -0.0034 | -0.0013 | -0.0309 | -0.1920 | 0.0468 | -0.0005 |
| Atg3 | 0.2665 | 0.1932 | -0.4966 | -0.3758 | 0.0166 | -0.0021 |
| COG3 | 0.0062 | 0.0009 | -0.0699 | 0.0236 | 0.0351 | -0.0329 |
| PD-1 | -0.0837 | -0.0773 | 0.0058 | 0.0952 | -0.0037 | 0.0379 |
| AMPKa_pT172 | 0.1530 | -0.0150 | 0.0008 | -0.2228 | -0.0573 | 0.1373 |
| PTEN | 0.0662 | 0.1081 | -0.0257 | -0.2147 | 0.0279 | -0.3274 |
| Gli1 | 0.0122 | -0.0525 | -0.0398 | 0.0517 | -0.0037 | 0.0182 |
| Bcl2 | -0.1832 | 0.0440 | 0.1510 | 1.3765 | -0.4504 | -0.0458 |
| TRIP13 | 0.0218 | 0.0835 | 0.1248 | -0.9378 | -0.0212 | -0.2044 |
| BAP1 | 0.0985 | -0.0739 | -1.8195 | -0.1835 | 0.0798 | 0.2354 |
| 14-3-3-beta | 0.0560 | 0.0161 | -0.0304 | 0.0851 | -0.1471 | -0.1300 |
| BiP-GRP78 | -0.0042 | -0.1031 | -0.0154 | -0.6449 | 0.0291 | 0.1284 |
| CSK | -0.0353 | 0.0195 | 0.0860 | -0.0445 | -0.0543 | 0.1325 |
| MCT4 | 0.1079 | 0.3316 | -0.0194 | -0.0691 | 0.0215 | -0.0896 |
| RPA32 | 0.3342 | -0.0278 | -0.0345 | -0.1397 | 0.0337 | 0.1638 |
| ER-a_pS118 | 0.0766 | -0.0279 | -0.0429 | 0.0524 | -0.0341 | 0.1541 |

| | | | | | | |
|---|---|---|---|---|---|---|
| Axl | 0.1392 | 0.0742 | -0.1607 | -0.0292 | 0.0701 | -0.1974 |
| Jak2 | -0.0263 | -0.1617 | -0.1192 | 0.3165 | 0.0269 | 0.1471 |
| GSK-3a-b | -0.0532 | -0.0666 | -0.0079 | 0.1694 | 0.0100 | 0.0253 |
| eEF2 | -0.1130 | 0.9607 | 0.0503 | 0.0544 | -0.0135 | -0.2713 |
| Claudin-7 | 0.0447 | 0.0224 | 0.0197 | -0.0231 | -0.0164 | -0.0924 |
| PAI-1 | 0.2077 | 0.2309 | -0.0414 | 0.0621 | -0.0738 | -0.2548 |
| PAK4 | 0.1164 | -0.0741 | 0.0038 | 0.0169 | -0.0033 | 0.2975 |
| Tuberin | 0.0586 | -0.0744 | -0.1062 | 0.4417 | -0.3168 | 0.1439 |
| DRP1 | 0.2193 | 0.0876 | -0.3068 | 0.0286 | 0.0122 | -0.0163 |
| Histone-H3_pS10 | -0.0803 | 0.2473 | 0.3023 | 0.0338 | 0.0070 | -0.0225 |
| Coup-TFII | 0.0073 | 0.0020 | -0.0163 | -0.2408 | 0.1539 | -0.1806 |
| GCLM | 0.0048 | -0.0565 | 0.0834 | 0.0143 | -0.0350 | 0.0638 |
| Smad4 | -0.0645 | 0.0319 | 0.0279 | -0.0088 | 0.0078 | 0.0067 |
| DAPK2 | -0.0270 | -0.0001 | -0.0196 | 0.1103 | 0.0060 | 0.0215 |
| Atg5 | 0.0296 | 0.0387 | -0.0493 | -0.1292 | 0.2596 | -0.2255 |
| Fibronectin | 0.0277 | -0.0812 | 0.0284 | 0.1112 | -0.0988 | -0.0341 |
| Mitofusin-1 | -0.0189 | 0.0998 | 0.0331 | 0.0221 | 0.0188 | -0.1298 |
| Cyclophilin-F | 0.1396 | -0.0049 | -0.8465 | -0.5463 | 0.0109 | 0.1051 |
| IGFBP3 | 0.0788 | -0.0708 | 0.0566 | -0.3782 | 0.1662 | -0.5491 |
| NRF2 | 0.0330 | 0.0675 | -0.0527 | -0.2026 | 0.1847 | -0.0450 |
| IGF1R_pY1135_Y1136 | -0.0153 | -0.0005 | -0.0350 | -0.5268 | 0.0462 | 0.0237 |
| STING | 0.1116 | 0.0587 | 0.0143 | -0.2829 | -0.0122 | -0.0495 |
| FAK_pY397 | 0.0073 | 0.0186 | 0.0355 | -0.6405 | -0.0890 | -0.0145 |
| MelanA | -0.0382 | 0.0193 | 0.1278 | -0.7508 | -0.0133 | 0.0450 |
| Heregulin | 0.1044 | -0.0368 | -0.0677 | -1.3042 | 0.1000 | 0.0350 |
| C-Raf | -0.0503 | 0.0289 | 0.0025 | -0.1951 | -0.0004 | 0.1192 |
| PR | 0.0002 | -0.0177 | 0.0274 | -0.2129 | 0.0003 | 0.0257 |
| TIGAR | 0.0149 | -0.0406 | -0.0116 | 0.1170 | -0.0500 | 0.0059 |
| b-Actin | 0.3518 | 0.2282 | 0.0734 | -0.0527 | -0.1137 | -0.2408 |
| Chk2_pT68 | -0.0011 | -0.1100 | -0.0749 | 0.2143 | 0.0016 | 0.1137 |
| SFRP1 | 0.0308 | 0.0160 | -0.0135 | -0.3241 | 0.0156 | -0.4906 |
| IL-6 | -0.0540 | -0.0658 | -0.0069 | 0.1984 | 0.0090 | 0.0532 |
| cGAS | -0.0059 | 0.0066 | 0.0167 | -0.2354 | -0.0030 | -0.0013 |
| Mnk1 | 0.1287 | -0.0401 | -0.0128 | 0.2808 | -0.0155 | 0.0071 |
| Smad3 | -0.0615 | 0.0115 | -0.2595 | 0.3308 | 0.1288 | -0.0133 |
| Bid | 0.0353 | -0.0025 | -0.0360 | 0.0229 | 0.0084 | 0.0132 |
| PRMT5 | -0.0537 | -0.0307 | -0.0333 | 0.1451 | 0.0354 | 0.0645 |
| 14-3-3-zeta | 0.3488 | 0.3477 | -0.7044 | -0.2805 | 0.3214 | -0.7006 |
| IRS2 | 0.0670 | -0.0273 | 0.1242 | -0.2034 | -0.3002 | 0.0255 |
| Rad17_pS645 | -0.0160 | 0.0868 | 0.0106 | -0.3115 | -0.0732 | 0.0088 |
| Mcl-1 | -0.0366 | -0.1388 | -0.0925 | 0.4336 | 0.0371 | 0.1784 |
| MMP14 | 0.0605 | 0.1945 | 0.0042 | -0.5023 | -0.0021 | -0.0603 |
| Notch1-cleaved | 0.0053 | 0.0023 | 0.0054 | -0.1111 | 0.0036 | -0.2204 |
| Twist | -0.0078 | -0.0418 | -0.0305 | 0.0923 | 0.0084 | 0.0489 |
| EphA2 | -0.0532 | 0.8701 | -0.3297 | -0.1446 | 0.5257 | 0.0460 |
| Stat3 | -0.0601 | 0.0733 | -0.1031 | 0.1017 | -0.1025 | 0.0529 |
| C-Raf_pS338 | 0.0050 | -0.1042 | -0.0749 | 0.5612 | -0.0045 | 0.2807 |
| FGFR2 | 0.0228 | -0.0540 | -0.0424 | 0.2828 | -0.0391 | 0.0996 |
| Atg4B | 0.0361 | 0.0136 | -0.0750 | 0.0567 | -0.0077 | -0.0434 |
| PEA-15 | -1.0215 | -0.5027 | -0.4005 | 0.7556 | 0.4026 | 0.5210 |
| PAX8 | -0.0018 | 0.0805 | 0.2602 | -0.2989 | -0.0682 | -0.0054 |

| | | | | | | |
|---|---|---|---|---|---|---|
| DNMT1 | 0.1193 | 0.0676 | 0.1528 | -0.7223 | -0.0617 | -0.2428 |
| FOXM1 | 0.1142 | 0.1257 | 0.1402 | -1.1612 | -0.1136 | -0.3862 |
| CD49b | -0.0202 | 0.0044 | 0.3648 | 0.1775 | -0.1197 | -0.1019 |
| GGPS1 | -0.0060 | -0.0755 | -0.0463 | 0.1271 | 0.0065 | 0.0575 |
| GAPDH | 0.2412 | 0.3931 | -0.3584 | 0.6943 | -0.2406 | -0.2732 |
| MAPK_pT202_Y204 | 0.2749 | 0.0036 | -0.0179 | -1.0350 | -0.0902 | 0.0736 |
| SGK3 | 0.1201 | 0.0052 | 0.0536 | -0.2493 | -0.0054 | -0.0070 |
| NQO1 | 0.0884 | -0.1042 | -0.2969 | 0.4315 | -0.5181 | 0.2129 |
| PERK | -0.0141 | 0.0223 | 0.3006 | -0.2321 | 0.0059 | 0.0069 |
| Calnexin | 0.1096 | -0.0711 | 0.0317 | -0.2849 | -0.0296 | 0.1373 |
| FABP5 | 0.1071 | 0.0528 | -0.0670 | -0.1491 | -0.5957 | 0.1341 |
| Rab25 | -0.1430 | -0.0143 | 0.0985 | 0.0657 | 0.0202 | -0.1824 |
| Syk | -0.5688 | 0.5530 | 1.6875 | -3.9118 | 1.8874 | -3.4896 |
| DDB-1 | -0.0355 | 0.0197 | -0.0488 | -0.1402 | 0.0903 | 0.0882 |
| SCD | 0.0167 | -0.1201 | -0.0363 | 0.5694 | -0.0424 | 0.2174 |
| SHP-2_pY542 | -0.0191 | 0.1205 | 0.1374 | -0.2550 | 0.0196 | -0.0437 |
| Cyclin-E1 | 0.2889 | -0.0770 | 0.2010 | -0.4959 | -0.2344 | 0.0752 |
| VHL | 0.1984 | 0.1309 | 0.0380 | -0.0404 | -0.0359 | -0.1753 |
| CRABP2 | -0.0370 | -0.0606 | 0.0717 | -0.4621 | 0.0583 | 0.0298 |
| CD68 | -0.0297 | -0.0537 | 0.0101 | -0.1486 | 0.0564 | 0.0245 |
| WTAP | -0.0887 | -0.2422 | -0.1929 | 0.1078 | 0.1608 | 0.2167 |
| Enolase-2 | 0.1922 | 0.1709 | 0.0272 | -0.0131 | -0.0251 | -0.0695 |
| GSK-3a-b_pS21_S9 | 0.0664 | 0.1508 | -0.0861 | 0.1398 | -0.1289 | -0.2012 |
| DUSP4 | 0.1619 | 0.7664 | 0.7500 | -0.1428 | -0.2749 | -0.4980 |
| GCLC | -0.0022 | 0.1873 | -0.1586 | 0.0213 | 0.1557 | -0.1324 |
| XPA | 0.0850 | -0.0406 | 0.0263 | -0.2065 | -0.0608 | 0.1751 |
| p38-MAPK | 0.3313 | 0.1787 | -0.0845 | -0.0435 | 0.0843 | -0.1842 |
| Raptor | 0.0203 | 0.0553 | 0.1005 | -0.1912 | -0.0198 | -0.0291 |
| GRB2 | 0.0857 | 0.0551 | 0.1464 | -0.1263 | -0.0491 | -0.2811 |
| Stat3_pY705 | -0.0073 | -0.0644 | -0.0124 | 0.1211 | -0.0921 | 0.0386 |
| Tyro3 | -0.0213 | -0.0126 | -0.0559 | 0.0651 | 0.0185 | 0.0679 |
| PAX6 | -0.0261 | -0.2346 | 0.0750 | 0.3774 | -0.1832 | 0.0189 |
| Lck | 0.0073 | 0.1249 | -0.2632 | 0.0771 | -0.0068 | -0.1785 |
| HER3 | -0.3151 | -0.4691 | 0.4756 | -2.0797 | 0.3156 | 0.7966 |
| HNRNPK | 0.0186 | 0.0041 | 0.1416 | -0.2017 | -0.0229 | -0.0059 |
| GSK-3B | 0.2354 | 0.0816 | 0.0505 | -0.1885 | -0.0484 | -0.1421 |
| Smad1 | -0.0452 | -0.1312 | -0.0073 | 0.0524 | 0.0094 | 0.1441 |
| PLC-gamma1_pS1248 | 0.0464 | 0.0260 | -0.0395 | -0.0509 | -0.0700 | 0.0339 |
| MSH2 | 0.0527 | -0.0004 | -0.1173 | -0.2345 | 0.0063 | 0.0476 |
| GRB7 | 0.0031 | -0.0191 | 0.1072 | 0.0160 | -0.0234 | 0.0546 |
| CD29 | 0.0335 | -0.0104 | -0.0038 | -0.1204 | 0.0451 | -0.2679 |
| PARG | -0.0321 | 0.0342 | 0.1802 | -0.0106 | -0.0962 | 0.0249 |
| 4E-BP1_pT37_T46 | 0.2002 | 0.0042 | -0.0184 | -0.0292 | -0.0376 | 0.0818 |
| MRAP | -0.0146 | 0.0443 | -0.0637 | 0.1088 | 0.0151 | -0.0235 |
| Ezh2 | -0.0456 | 0.6838 | 0.0259 | -0.4001 | 0.0690 | -0.6904 |
| MDM2_pS166 | -0.0110 | 0.0284 | 0.0142 | -0.5369 | 0.0115 | -0.1294 |
| Oct-4 | -0.1187 | 0.1214 | -0.1184 | 0.2444 | 0.0025 | 0.0120 |
| RIP3 | -0.0323 | -0.0208 | -0.0513 | 0.1248 | 0.0298 | 0.0190 |
| Gys_pS641 | -0.0606 | -0.0488 | -0.0235 | 0.1519 | 0.0256 | 0.0759 |
| LC3A-B | -0.2782 | 0.3950 | 0.1482 | -0.1275 | 0.1889 | -0.2330 |
| MMP2 | -0.0304 | 0.0146 | 0.0306 | -0.0205 | 0.0335 | -0.0215 |

| | | | | | | |
|---|---|---|---|---|---|---|
| DDR1 | 0.0164 | -0.0387 | 0.2171 | -0.3523 | 0.0259 | -0.0114 |
| PI3K-p85 | 0.0137 | 0.0530 | 0.0284 | -0.9903 | -0.0197 | -0.0209 |
| FTO | 0.0279 | -0.0347 | -0.0185 | 0.1702 | -0.1748 | 0.0129 |
| Complex-II-Subunit | 0.0074 | -0.0020 | -0.0054 | -0.2959 | 0.0080 | -0.1063 |
| EVI1 | -0.0048 | -0.0558 | 0.0564 | 0.0239 | -0.0592 | 0.0267 |
| Vinculin | 0.5277 | 0.4016 | 0.4548 | -0.3771 | -0.7606 | -0.5501 |
| Aurora-B | -0.1763 | 0.1836 | 0.1567 | -0.3698 | 0.2102 | -0.7949 |
| MERIT40_pS29 | -0.0350 | 0.0440 | 0.0746 | 0.0541 | -0.0349 | -0.0384 |
| PRMT1 | -0.0093 | 0.3827 | -0.0495 | -0.0808 | 0.0098 | 0.1154 |
| Tau | -0.2527 | -0.3033 | 0.2618 | 0.3371 | -0.0793 | 0.0938 |
| NAPSIN-A | 0.0140 | -0.0298 | 0.0543 | -0.0712 | -0.0433 | 0.0932 |
| MITF | 0.1214 | -0.0545 | -0.0552 | 0.3379 | -0.1639 | 0.0496 |
| Akt2_pS474 | -0.0225 | 0.3238 | 0.4201 | -0.7184 | 0.0231 | -0.1632 |
| Akt1_pS473 | -0.0164 | -0.0105 | -0.0559 | 0.0577 | 0.0263 | 0.0087 |
| DM-Histone-H3 | 0.0224 | 0.0708 | -0.0218 | -0.0790 | 0.0239 | -0.1910 |
| Cox-IV | 0.1075 | 0.2445 | 0.5730 | -0.5094 | -0.1070 | -0.3455 |
| Glucocorticoid-Receptor | -0.0832 | 0.7690 | 0.7737 | 0.1023 | -0.3573 | -0.1594 |
| FAK | -0.0468 | 0.0581 | -0.1240 | 0.1157 | -0.0332 | 0.0396 |
| CD38 | 0.0160 | -0.0318 | 0.0178 | -0.2608 | -0.0614 | 0.0751 |
| AR | -0.0174 | -0.0130 | 0.0681 | -0.1852 | 0.0179 | 0.0399 |
| ADAR1 | 0.0229 | -0.0031 | 0.0262 | -0.3596 | 0.0091 | -0.2298 |
| Transglutaminase | 0.4025 | 0.3294 | -0.4928 | 0.3427 | -0.5097 | -0.3096 |
| Lasu1 | -0.0366 | -0.2060 | -0.3257 | 0.7901 | 0.0372 | 0.2294 |
| DNA_POLG | 0.1046 | -0.0280 | -0.0108 | 0.1399 | -0.0510 | 0.0052 |
| RSK1 | 0.1111 | 0.1219 | 0.2234 | -0.2920 | -0.1106 | -0.6074 |
| PMS2 | 0.0308 | -0.0115 | -0.0028 | -0.2860 | -0.0064 | 0.1334 |
| MACC1 | -0.0103 | -0.0575 | 0.2356 | -0.4816 | 0.0108 | 0.0634 |
| Pyk2_pY402 | 0.0429 | 0.0699 | -0.0432 | -0.6500 | 0.0453 | -0.3068 |
| p38-MAPK-_pT180_Y182 | 0.0044 | -0.3768 | 0.2140 | -0.0428 | -0.0038 | 0.3065 |
| S6_pS240_S244 | 0.0320 | 0.1345 | 0.0702 | -0.6088 | -0.0314 | -0.4002 |
| p27_pT198 | -0.0651 | -0.0519 | -0.0020 | 0.1769 | 0.0041 | 0.1052 |
| Collagen-VI | -0.0124 | -0.0348 | -0.0072 | 0.2836 | -0.0271 | 0.1357 |
| CRABP1 | 0.0007 | -0.0165 | 0.1873 | -0.0786 | 0.1396 | -0.0651 |
| PAICS | -0.8298 | -0.0029 | -0.6953 | 0.2431 | 0.0089 | 0.2504 |
| mTOR_pS2448 | 0.0379 | 0.0465 | -0.1240 | 0.2050 | -0.0374 | -0.6119 |
| INPP4b | 0.1579 | 0.0246 | 0.0510 | -0.3914 | -0.0187 | -0.3655 |
| Merlin | 0.0261 | 0.2127 | -0.0458 | 0.1368 | -0.0478 | -0.1045 |
| Gli3 | -0.0957 | 0.0060 | 0.0064 | -0.0236 | 0.0103 | 0.0042 |
| CD134 | -0.0155 | -0.0023 | 0.0256 | -0.0113 | 0.0169 | 0.0005 |
| HLA-DR-DP-DQ-DX | 0.0950 | 0.0430 | -0.0773 | -0.0479 | -0.0347 | 0.0493 |
| B-Raf_pS445 | -0.0016 | 0.0056 | -0.0180 | -0.0182 | 0.0217 | -0.0116 |
| N-Ras | -0.0062 | -0.0479 | -0.0135 | 0.2989 | -0.0547 | 0.0336 |
| p70-S6K1 | 0.0078 | 0.0891 | -0.0684 | 0.1079 | -0.0073 | -0.0727 |
| ENY2 | 0.0373 | -0.1584 | -0.0570 | 0.6642 | -0.0782 | 0.3535 |
| XPF | -0.0758 | 0.0144 | 0.1119 | 0.0102 | 0.0969 | -0.0223 |
| Rheb | -0.0850 | 1.1239 | -1.2472 | 0.8583 | -0.1055 | 0.0778 |
| Connexin-43 | 0.0097 | 0.1075 | 0.0851 | -0.6822 | -0.0092 | -0.7351 |
| PKCa | -0.0133 | 0.1197 | 0.1295 | 0.0325 | -0.2946 | -0.2438 |
| Gys | -0.2314 | -0.0802 | -0.0156 | 0.1221 | 0.0626 | 0.0100 |
| AceCS1 | 0.2616 | 0.1671 | 0.0174 | 0.0033 | -0.0754 | -0.0942 |
| PKM2 | 0.1019 | 0.4765 | -0.1216 | -0.1148 | 0.1929 | -0.1252 |

| | | | | | | |
|---|---|---|---|---|---|---|
| p38-a | 0.1037 | 0.0031 | -0.3283 | 0.1022 | -0.0986 | -0.0049 |
| ACSL1 | -0.1379 | -0.0258 | -0.1453 | 0.4677 | 0.0318 | 0.2622 |
| Aurora-A | 0.1782 | 0.1990 | 0.1625 | -0.4226 | -0.1604 | -0.5422 |
| VASP | 0.1474 | -0.0865 | 0.0723 | -0.8726 | -0.2855 | 0.1291 |
| CDK1_pT14 | 0.4660 | 0.3677 | 0.2889 | -1.3851 | -0.2868 | -0.7894 |
| ACLY_pS455 | -0.0097 | 0.1439 | -0.0100 | -0.0462 | -0.0392 | 0.0054 |
| FOXO3 | 0.0342 | 0.0089 | -0.0168 | 0.0374 | -0.0250 | -0.0095 |
| SOX7 | -0.0111 | -0.0047 | -0.0423 | 0.2279 | -0.0123 | 0.0395 |
| XIAP | 0.0040 | -0.0742 | -0.1197 | 0.2120 | -0.0035 | 0.0813 |
| HMHA1 | -1.1001 | 1.6548 | -1.1300 | 0.1632 | 1.1296 | -0.1301 |
| AMPKa | -0.0111 | 0.0488 | -0.0815 | 0.1410 | -0.0349 | 0.0039 |
| VEGFR2_pY1175 | 0.0114 | 0.0314 | 0.0746 | -0.4220 | -0.0642 | -0.0187 |
| FRS2-alpha_pY196 | -0.0362 | -0.0260 | 0.0226 | -0.3948 | 0.2280 | 0.0242 |
| Stathmin-1 | 0.0187 | -0.1981 | -0.1390 | 0.3307 | -0.0182 | 0.0725 |
| Cox2 | 0.0043 | -0.0067 | -0.0359 | 0.1540 | 0.0085 | 0.0048 |
| Akt1 | 0.0633 | 0.1284 | 0.0238 | -0.0031 | -0.0552 | -0.0599 |
| RRM2 | -0.0133 | 0.3313 | -0.0113 | 0.0450 | 0.0134 | -0.1907 |
| SHP2 | 0.0117 | -0.0275 | 0.0855 | 0.1912 | -0.0915 | -0.0195 |
| PAK1 | -0.0197 | -0.1284 | 0.5322 | 0.0388 | 0.1607 | -0.0918 |
| Cyclin-B1 | 0.0767 | -0.0131 | 0.5376 | -0.2297 | -0.3069 | 0.0113 |
| GATA3 | -0.1494 | 0.2692 | -0.2634 | 0.4646 | -0.0353 | 0.0498 |
| TTF1 | 0.0571 | 0.1408 | -0.0768 | 0.1808 | -0.0974 | -0.0939 |
| p90RSK_pT573 | 0.2387 | -0.0083 | 0.0020 | -2.3897 | -0.2157 | 0.0065 |
| DAPK1_pS308 | 0.0645 | 0.2653 | -0.4366 | 0.1067 | -0.0639 | -0.0791 |
| ATP5A | -0.8120 | 0.0711 | 0.0510 | 0.1692 | -0.3321 | -0.0566 |
| b-Catenin | -0.5199 | 0.0749 | -0.2869 | 1.0585 | -0.0690 | 0.1671 |
| ACC1 | 0.0076 | -0.0234 | -0.5063 | 0.1836 | -0.0955 | 0.0575 |
| KAP1 | 0.1267 | -0.0413 | 0.0270 | -0.2161 | 0.0490 | -0.3192 |
| TUFM | 0.0649 | -0.0021 | -0.1234 | 0.0266 | 0.0442 | -0.1643 |
| PLC-gamma2_pY759 | -0.0187 | 0.0473 | 0.2702 | 0.0378 | -0.0985 | -0.0108 |
| IMP3 | -0.0129 | -0.0029 | 0.0131 | 0.0234 | -0.0019 | 0.0874 |
| Rictor_pT1135 | 0.0201 | 0.0300 | 0.0312 | -0.1961 | -0.0862 | -0.0274 |
| Bak | -0.0121 | -0.0301 | 0.1251 | 0.0256 | 0.0126 | 0.1874 |
| mTOR | 0.1146 | -0.0310 | -0.4031 | 0.0555 | -0.0319 | 0.0471 |
| JNK_pT183_Y185 | -0.0053 | -0.0397 | -0.0143 | 0.3861 | -0.0214 | 0.1033 |
| ULK1_pS757 | -0.0278 | 0.0120 | 0.0267 | -0.0767 | 0.0594 | -0.1441 |
| eIF4E | 0.0144 | 0.2159 | -0.0341 | -0.2260 | 0.1231 | -0.3725 |
| Chk2 | -0.0269 | -0.0180 | 0.0317 | -0.1588 | 0.0239 | 0.2532 |
| UBQLN4 | -0.0226 | -0.0657 | 0.0027 | 0.1079 | -0.0006 | 0.0660 |
| MR1 | -0.0458 | -0.0790 | -0.0028 | 0.0871 | 0.0049 | 0.0692 |
| PYGM | -0.0175 | 0.0017 | -0.0946 | 0.2648 | -0.0003 | 0.0455 |
| Rad51 | -0.0163 | -0.0121 | 0.0223 | 0.0047 | 0.0168 | 0.0337 |
| D-a-Tubulin | -0.0147 | -0.0145 | 0.0011 | 0.0338 | 0.0283 | -0.0349 |
| EGFR | 0.1189 | 0.1009 | -0.1157 | 0.1460 | -0.1432 | -0.1027 |
| Chk1 | -0.0299 | 0.0188 | 0.0913 | 0.0490 | -0.0156 | -0.0181 |
| CD45 | -0.1409 | -0.1505 | -0.0413 | 0.1034 | 0.0434 | 0.1089 |
| S6 | -0.0139 | 0.3276 | -0.0926 | 0.0004 | 0.6326 | 0.0067 |
| YB1_pS102 | 0.0294 | -0.0656 | -0.1237 | 0.3496 | -0.0289 | 0.0931 |
| Gab2 | 0.0730 | -0.0029 | -0.0113 | -0.1380 | -0.0010 | 0.1745 |
| PDHK1 | -0.0096 | -0.0023 | -0.0523 | -0.1013 | 0.0101 | 0.0087 |
| Glutaminase | 0.0296 | -0.0454 | -0.2837 | 0.1457 | 0.1380 | -0.1254 |

| | | | | | | |
|---|---|---|---|---|---|---|
| eEF2K | 0.0900 | 0.3760 | -0.0750 | 0.0957 | -0.3289 | -0.1448 |
| Bax | 0.0248 | -0.1268 | 0.0090 | 0.1876 | -0.0069 | -0.1046 |
| UGT1A | -0.0394 | 0.0236 | 0.1722 | -0.7808 | 0.0995 | -0.8174 |
| LDHA | 0.1254 | -0.0318 | 0.0860 | -0.6117 | -0.0253 | 0.0300 |
| Dvl3 | -0.0875 | -0.0194 | -0.0410 | 0.3789 | 0.0253 | 0.0798 |
| CDT1 | -0.0189 | -0.0798 | -0.0007 | 0.2632 | -0.0922 | 0.0655 |
| Myosin-IIa_pS1943 | -0.0626 | 0.1305 | -0.2696 | 0.1997 | 0.0632 | -0.1030 |
| CD74 | -0.0034 | 0.0652 | 0.2226 | -0.2264 | -0.0163 | -0.0038 |
| Bim | -1.2975 | -0.0031 | 0.9059 | -1.3106 | 0.0091 | 0.2520 |
| Smac | -0.6122 | 0.2641 | -0.5022 | -0.1359 | 0.1767 | 0.4330 |
| DNA-Ligase-IV | -0.0145 | -0.0895 | -0.0051 | 0.0487 | -0.0665 | 0.0742 |
| MTCO1 | -0.3400 | 0.3989 | -0.1124 | 1.0926 | 0.0596 | -0.0451 |
| p27_pT157 | -0.0090 | -0.0347 | -0.0749 | 0.1319 | 0.0095 | 0.0331 |
| SLC1A5 | 0.1315 | -0.0729 | -0.1864 | -0.0432 | 0.0788 | 0.4871 |
| Cadherin-6 | -0.0552 | -0.1290 | 0.0551 | 0.0385 | 0.1242 | -0.0054 |
| cdc25C | 0.0243 | -0.1322 | -0.0439 | 0.1081 | -0.0757 | 0.1426 |
| Enolase-1 | 0.6704 | 0.3543 | -0.3666 | 0.1667 | -0.1258 | -0.3118 |
| MERIT40 | 0.0797 | 0.0353 | -0.5511 | -0.3189 | -0.0293 | 0.0445 |
| PREX1 | -0.1001 | -0.1755 | 0.2800 | 0.3037 | -0.1853 | 0.0929 |
| Bad_pS112 | 0.0290 | -0.0950 | -0.1462 | 0.2752 | -0.0284 | 0.2117 |
| ER-a | -0.1622 | -0.1399 | -0.0005 | 0.6933 | 0.0026 | 0.2304 |
| PI3K-p110-b | -0.0147 | -0.0431 | -0.0099 | 0.2080 | 0.0120 | 0.0510 |
| FoxO3a_pS318_S321 | 0.0020 | -0.0178 | 0.0481 | -0.5688 | -0.0578 | 0.1101 |
| NDUFB4 | 0.0096 | 0.0069 | 0.0021 | -0.2525 | -0.0427 | -0.0077 |
| ASNS | -0.3168 | 0.4397 | 0.3268 | -0.3216 | 0.3174 | -0.3955 |
| cdc2_pY15 | 0.0575 | 0.1247 | -0.0004 | -0.1616 | 0.0025 | -0.6740 |
| AMPK-a2_pS345 | -0.0054 | -0.0133 | -0.0701 | 0.1781 | 0.0059 | 0.0630 |
| DYRK1B | -0.0157 | 0.0421 | 0.0541 | -0.0676 | 0.0162 | -0.1593 |
| Rad23A | -0.0210 | 0.0161 | -0.1166 | 0.1441 | -0.1670 | 0.0138 |
| BMK1-Erk5_pT218_Y220 | -0.0020 | -0.0660 | 0.0320 | -0.1434 | 0.0026 | 0.1801 |
| TRIM25 | -0.0379 | 0.0223 | 0.0182 | -0.5395 | 0.1574 | -0.0652 |
| PARP | 0.0008 | -0.0166 | 0.1940 | -0.5555 | -0.0590 | 0.2146 |
| 4E-BP1_pS65 | 0.1083 | -0.1141 | -0.6053 | 0.2037 | -0.1433 | 0.1122 |
| FGFR1 | -0.1246 | -0.0916 | 0.0801 | 0.0993 | 0.0818 | -0.0662 |
| Bcl-xL | 0.0017 | -0.0502 | -0.1790 | 0.0576 | 0.0657 | -0.0090 |
| VHL-EPPK1 | -0.1886 | 2.1278 | -0.2676 | 0.1261 | 0.1749 | -0.0930 |
| Paxillin | 0.1682 | 0.0961 | 0.0348 | -0.4202 | -0.0327 | -0.2939 |
| Beclin | -0.0235 | -0.0557 | 0.0457 | 0.0194 | 0.0215 | 0.1198 |
| Chk1_pS345 | -0.0924 | -0.1108 | -0.0400 | 0.3494 | 0.0421 | 0.0646 |
| ACC_pS79 | -0.0153 | 0.6961 | 0.4240 | -0.0211 | 0.0158 | -0.3947 |
| HSP27_pS82 | 0.3086 | 0.3347 | 0.0031 | -0.3238 | -0.0010 | -0.4187 |
| H2AX_pS140 | -0.0098 | -0.0060 | -0.1256 | 0.0748 | 0.0399 | -0.0152 |
| 53BP1 | -0.0819 | -0.1407 | -0.3658 | 0.4038 | 0.0825 | 0.2870 |
| Creb | -0.0154 | -0.0400 | -0.0518 | 0.1564 | 0.0297 | 0.0081 |
| Midkine | 0.0207 | -0.0365 | 0.1092 | -0.1321 | -0.0227 | 0.0379 |
| Sox2 | 0.1987 | -0.9342 | 0.1150 | -0.0943 | 0.5998 | -0.5006 |
| IGFRb | -0.1341 | -0.1060 | -0.3917 | 0.5663 | 0.1120 | 0.2825 |
| 14-3-3-epsilon | -0.0095 | -0.0625 | -0.0102 | 0.4285 | -0.0021 | 0.1068 |
| Chk1_pS296 | -0.0680 | 0.0338 | -0.0347 | 0.0554 | 0.1384 | -0.1035 |
| PCNA | 0.0570 | -0.0332 | 0.0189 | -0.0355 | -0.0315 | 0.1758 |
| EGFR_pY1173 | 0.0286 | 0.0256 | -0.5075 | 0.4226 | -0.0197 | -0.0971 |

| | | | | | | |
|---|---|---|---|---|---|---|
| Rb | -0.1418 | 0.0001 | 0.0695 | -0.1624 | 0.3299 | -0.0019 |
| MLH1 | -0.0571 | -0.0200 | -0.1559 | 0.2257 | 0.5959 | 0.0182 |
| Caspase-3-cleaved | -0.0204 | -0.0501 | -0.0273 | 0.2330 | 0.0210 | 0.0895 |
| PEA-15_pS116 | 0.0115 | -0.0476 | -0.0093 | -0.0359 | 0.0114 | 0.0803 |
| Atg7 | 0.0796 | -0.0358 | -0.1748 | 0.0964 | -0.1950 | 0.0340 |
| YES1 | 0.0038 | -0.0467 | 0.1012 | -0.4043 | -0.0033 | 0.0902 |
| Pdcd4 | -0.1687 | -0.4200 | -0.1216 | 1.2153 | 0.1237 | 0.3282 |
| XBP-1 | -0.1197 | -0.4749 | -0.5103 | 0.1881 | 0.3775 | 0.1125 |
| MIF | -0.1813 | 0.1798 | 0.1816 | 0.0978 | -0.1522 | -0.0647 |
| Aurora-ABC_pT288_pT232_pT198 | 0.0262 | -0.0420 | 0.0741 | -0.4795 | -0.1275 | 0.0964 |
| VEGFR-2 | -0.4125 | 0.2247 | -0.0606 | 0.0813 | 0.1341 | -0.2537 |
| Snail | -0.0065 | -0.0093 | 0.6123 | -0.6617 | 0.2583 | -0.1991 |
| Histone-H3 | -0.1932 | 0.3171 | 0.1736 | -0.9741 | 0.2216 | -0.2067 |
| UQCRC2 | -0.0624 | -0.0565 | 0.0532 | 0.3060 | -0.0277 | 0.0422 |
| PHLPP | 0.1421 | 0.0263 | -0.0029 | -0.3035 | 0.0050 | -0.1462 |
| b-Catenin_pT41_S45 | -0.0172 | -0.0823 | -0.0621 | 0.3542 | 0.0177 | 0.1341 |
| FGF-basic | 0.0875 | 0.1089 | -0.0421 | -0.0607 | -0.2362 | 0.0364 |
| Granzyme-B | -0.2866 | 0.1403 | 0.0254 | 0.0764 | -0.0233 | -0.2279 |
| PIP4K2A | 0.0527 | -0.0320 | -0.0662 | 0.0184 | 0.0225 | 0.0584 |
| BCL2A1 | -1.7994 | -0.0817 | -0.2381 | 0.4442 | 0.0877 | 0.0987 |
| HER3_pY1289 | -0.0152 | -0.0029 | -0.1550 | 0.5170 | 0.0089 | 0.1474 |
| PDHA1 | -0.0281 | -0.1136 | -0.0161 | 0.4081 | 0.0182 | 0.1854 |
| PI3K-p110-a | -0.0406 | -0.0804 | -0.0384 | 0.3484 | 0.0405 | 0.0996 |
| CD2 | -0.0558 | -0.1128 | 0.0099 | 0.1680 | -0.0078 | 0.0416 |
| 4E-BP1 | -0.2197 | -0.0553 | -0.2569 | 0.3046 | 0.0612 | 0.2308 |
| ATP5H | 0.1435 | 0.0215 | -0.0358 | -0.3234 | -0.0517 | 0.0355 |
| Jagged1 | 0.0111 | 0.0690 | -0.0315 | -0.0573 | 0.0953 | -0.0183 |
| VAV1 | -0.2069 | -0.1585 | 0.0643 | 0.5362 | -0.0622 | 0.1551 |
| Hif-1-alpha | -0.0753 | -0.0930 | 0.0150 | 0.0840 | 0.0556 | -0.0207 |
| HSP60 | -0.0180 | -0.0358 | 0.0077 | 0.2544 | 0.0185 | -0.1260 |
| PKC-b-II_pS660 | 0.2346 | 0.2097 | 0.1428 | -0.3040 | -0.1530 | -0.1485 |
| TFAM | -0.0849 | -0.1995 | 0.3643 | 0.5247 | -0.2623 | 0.0777 |
| c-Myc | -0.0142 | -0.1507 | -0.0054 | -1.1711 | 0.1055 | 0.2975 |
| Caspase-7-cleaved | -0.0566 | -0.1162 | -0.0234 | 0.2599 | 0.0255 | 0.1178 |
| Src_pY416 | 0.3185 | 0.2471 | 0.3805 | -5.4206 | -0.2411 | -1.7142 |
| MTSS1 | -0.0606 | -0.0863 | 0.0198 | 0.1313 | -0.0177 | 0.0967 |
| YAP_pS127 | 0.2055 | -0.0045 | -0.1320 | 0.0290 | -0.2976 | 0.0583 |
| PRC1_pT481 | -0.0495 | -0.0311 | 0.0459 | 0.2359 | -0.0252 | 0.0293 |
| Rad50 | 0.0276 | -0.0434 | -0.2208 | 0.1654 | -0.0402 | 0.0686 |
| WIPI1 | -0.0382 | 0.0517 | -0.2264 | 0.0598 | -0.0377 | 0.0309 |
| Ets-1 | -0.2521 | 0.0599 | 0.1638 | -0.0354 | 0.0802 | -0.2335 |
| IRF-3 | 0.0995 | 0.0090 | 0.0784 | -0.1508 | -0.0031 | -0.1129 |
| TRAP1 | 0.1347 | -0.0740 | -0.1880 | -0.1881 | 0.2140 | 0.0722 |
| Caveolin-1 | 0.0749 | -0.0873 | 0.8129 | -0.1112 | -0.3026 | 0.0855 |
| CHD1L | 0.0433 | 0.0171 | -0.0007 | -0.3049 | -0.0545 | -0.0049 |
| Hexokinase-I | -0.1038 | 0.1310 | 0.3777 | 0.0173 | 0.0043 | 0.0102 |
| Slfn11 | 0.0821 | -0.0590 | -0.3845 | 0.3889 | 0.0649 | -1.0710 |
| Elk1_pS383 | 0.0269 | -0.0301 | 0.0158 | 0.1584 | -0.0488 | -0.0524 |
| PLC-gamma1 | -0.3920 | 0.4680 | 0.0690 | -0.4796 | 0.4442 | -0.0746 |
| TROP2 | -0.0635 | 0.1452 | -0.0187 | 0.0394 | 0.1416 | -0.1024 |

| | | | | | | |
|---|---|---|---|---|---|---|
| HLA-DQA1 | -0.4305 | -0.0066 | -0.0076 | 0.0694 | 0.0397 | -0.0784 |
| RRM1 | -0.0245 | 0.0087 | 0.0534 | -0.0663 | -0.0150 | 0.0357 |
| Puma | -3.0366 | -0.0130 | 0.2896 | -0.5754 | 0.2850 | 0.0112 |
| MEK2 | -0.0030 | -0.0296 | -0.0250 | 0.1942 | 0.0036 | 0.0724 |
| Mitofusin-2 | -0.0443 | 0.0948 | 0.0197 | -0.1466 | -0.0176 | 0.1806 |
| CD171 | -1.2696 | -0.1365 | 0.1223 | 0.4556 | 0.2282 | -0.3567 |
| SF2 | -0.1324 | -0.0242 | 0.1904 | 0.0488 | 0.0890 | -0.1131 |
| B7-H4 | -0.0275 | 0.0308 | 0.0079 | -0.2230 | 0.0362 | -0.0684 |
| Akt_pT308 | 0.1476 | 0.2142 | 0.0012 | 0.0195 | -0.2649 | -0.2471 |
| PRAS40_pT246 | 0.0349 | -0.2039 | -0.3108 | 0.8744 | -0.0344 | 0.6522 |
| Notch3 | -0.0278 | -0.0201 | 0.0101 | 0.0304 | 0.0344 | 0.0027 |
| Annexin-I | 0.4316 | 0.1741 | -0.0418 | -0.1006 | 0.0439 | -0.4262 |
| ZAP-70 | -0.0144 | -0.0127 | 0.1227 | 0.0102 | 0.0445 | 0.0071 |
| Caspase-8 | 0.1216 | -0.0026 | -0.0116 | 0.1920 | -0.0284 | -0.0662 |
| GATA6 | -0.0035 | -0.0123 | 0.2500 | -0.2470 | -0.0334 | 0.1052 |
| N-Cadherin | -0.0174 | -0.0597 | 0.1275 | 0.0365 | -0.1074 | 0.0729 |
| PAK_pT423_T402 | 0.0275 | 0.0150 | -0.2765 | 0.1550 | -0.0091 | -0.5099 |
| Tuberin_pT1462 | 0.0367 | 0.0798 | 0.3517 | -0.8534 | -0.0362 | -0.8739 |
| YTHDF2 | 0.0131 | -0.0289 | 0.4029 | -0.5825 | -0.1160 | 0.0470 |
| Src | 0.0444 | -0.0053 | -0.2041 | -0.3705 | 0.0133 | 0.0035 |
| p53 | -0.0139 | 0.1243 | -0.0057 | -0.1057 | 0.0306 | -0.0912 |
| B7-H3 | -0.0239 | -0.0608 | -0.1739 | 0.0632 | 0.3369 | 0.0167 |
| CDK9 | 0.0529 | -0.0031 | 0.0233 | -0.1491 | -0.0566 | 0.0013 |
| IR-b | -0.1718 | -0.4279 | 0.3881 | 0.1322 | 0.4965 | -0.0991 |
| HER2 | -0.0369 | 0.0752 | -0.6789 | 0.3782 | 0.0374 | -0.2355 |
| Rab11 | 0.0088 | 0.0131 | 0.0049 | -0.0004 | -0.0028 | -0.0158 |
| PDH | -0.0048 | -0.0391 | -0.0388 | 0.1262 | 0.0054 | 0.0534 |
| Porin | -0.0617 | -0.0209 | 0.0672 | 0.0455 | -0.0199 | 0.0496 |
| METTL3 | -0.1073 | -0.0204 | -0.0376 | 0.0544 | 0.0264 | 0.0261 |
| PD-L1 | -0.0036 | -0.0122 | -0.1041 | 0.1759 | 0.0194 | -0.1352 |
| PRAS40 | -0.0477 | -0.0701 | 0.0238 | 0.0447 | -0.0038 | 0.0298 |
| Patched | -0.0186 | 0.0092 | -0.0353 | 0.0554 | 0.0192 | -0.0210 |
| HES1 | 0.0393 | 0.0325 | -0.0467 | -0.1388 | 0.1605 | -0.3697 |
| FN14 | 0.0264 | 0.1856 | -0.0461 | -0.5998 | 0.0702 | -0.0762 |
| JNK2 | -0.7032 | -0.1473 | -1.0520 | 0.1719 | 0.8380 | 0.4729 |
| CD5 | -0.0739 | -0.0929 | -0.0071 | 0.1228 | 0.0092 | 0.0732 |
| Ambra1_pS52 | -0.0797 | 0.0018 | -0.0161 | -0.1657 | 0.0294 | 0.0324 |
| XRCC1 | 0.0502 | -0.1879 | 0.1378 | 0.3891 | -0.4358 | -0.0575 |
| c-IAP2 | -0.0587 | 0.2901 | -0.1135 | -0.2429 | 0.0592 | 0.4091 |
| CDKN2A | 0.3621 | 0.3335 | 0.1987 | -1.6723 | -0.1966 | -1.1377 |
| Shc_pY317 | -0.5026 | -0.2236 | -0.5532 | 0.2481 | 1.0796 | 0.6385 |
| S6_pS235_S236 | 0.1767 | 0.1910 | 0.1205 | -0.3666 | -0.1184 | -0.2826 |
| U-Histone-H2B | -0.0122 | -0.0036 | -0.0367 | -0.0112 | 0.0226 | 0.0236 |
